# Immune competence and spleen size scale with colony status in the naked mole-rat

**DOI:** 10.1101/2021.08.15.456351

**Authors:** Valérie Bégay, Branko Cirovic, Alison J. Barker, Robert Klopfleisch, Daniel W. Hart, Nigel C. Bennett, Gary R. Lewin

## Abstract

Naked mole-rats (NM-R; *Heterocephalus glaber*) live in multi-generational colonies with a social hierarchy, show low cancer incidence and long life-spans. Here we asked if such extreme physiology might have an immune component. The spleen is the largest lymphoid organ and plays an essential role in response to immunological insults and may participate in combating cancer and slowing ageing. We investigated the anatomy, molecular composition and function of the NM-R spleen using RNA-sequencing and histological analysis in healthy animals. We found that spleen size in healthy NM-Rs varies considerably. We therefore classified NM-Rs according to spleen size as NM-Rs with small spleens or enlarged spleens. Animals with enlarged spleens showed potentially better anti-microbial profiles and were much more likely to have a high rank within the colony. Splenomegaly was associated with infection in sick NM-Rs, but not in NM-Rs with enlarged spleens. In all healthy NM-Rs splenic erythropoiesis, megakaryopoiesis and myelopoiesis were increased, but B lymphopoiesis was reduced and splenic marginal zone showed markedly altered morphology when compared to other rodents. However, in NM-Rs lymphocytes were found in secondary sites such as lymph nodes, gut lymphoid nodules and thymus. Thus, the NM-R spleen is a major site of adult hematopoiesis under normal physiological conditions. Overall, the NM-R immune system seems to rely mainly on innate immune responses with a more restricted adaptive immune response. We propose that the anatomical plasticity of the spleen might be regulated by social interaction and gives immunological advantage to increase the life-span of higher ranked animals.

## Introduction

Disease susceptibility is regulated by multiple factors including environmental stress and genetic factors. The immune system plays a critical role in protecting animals from infections, cancer and optimal immune function is associated with healthy ageing [1, 2]. In cases of pathogenic insult, the immune system protects the organism by engaging both innate and adaptive immune responses via either myeloid cells (granulocytes, macrophages and monocytes) and natural killer cells (NK) or lymphocytes and dendritic cells, respectively. Deregulation of the immune system is a critical factor in the development of cancer and ageing as immune function declines with age [2].

Naked mole-rats (NM-Rs; *Heterocephalus glaber*) show an extraordinarily long life-span for their small size (>30 years) [3, 4] and display a low cancer incidence [5–7]. Many features of NM-R physiology and habitat might contribute to the low cancer incidence, such as unique metabolic adaptations and hypoxia tolerance [8–10]. Recently, it has been shown that transformed NM-R cells can form tumours in mice [11], suggesting that non-cell autonomous mechanisms might eliminate tumorigenic cells before their spread in NM-Rs. Thus, the NM-R shows promise as an animal model to study the role of the immune system in cancer and ageing. NM-Rs are eusocial mammals that live in large colonies (on average 40-70 individuals), dominated by the queen who is normally the only breeding female [12, 13]. In our laboratory we have kept NM-R breeding colonies for more than 10 years. Over the last 4 years we have been monitoring the health status and mortality of our NM-Rs which only rarely die in captivity. Indeed, we observed only one major cause of death which was following fights with rivals during attempts to replace the breeding queen. Often wounded animals have unhealed infected wounds and have to be euthanized. *In vivo* experiments have shown that NM-Rs did not survive viral infections due to Coronavirus or Herpes simplex virus [14, 15]. Single-cell-RNA sequencing analysis of the spleen and of the peripheral blood of young adults showed that NM-Rs have a high myeloid to lymphoid cell ratio, but appear to lack classic NK cells [16]. These observations suggest that the NM-R immune system may differ significantly from that of conventional laboratory rodents.

In adulthood, secondary lymphoid organs like the spleen and lymph nodes participate in immune homeostasis. In humans and rodents extramedullary hematopoiesis takes place in the spleen to support adult bone marrow hematopoiesis under stress conditions [17, 18]. In addition, the spleen can also supply cells that stimulate cancer progression in mouse tumour models [19, 20]. Hence depending on the context, the spleen may support hematopoiesis, prevent growth of cancer cells or facilitate the development of tumour in mice. In NM-R little is known about the structure and function of the spleen in normal physiological and pathological conditions. Here we investigated the role of the spleen in healthy NM-Rs using molecular profiling and anatomical analysis.

We show that the size of the spleen varies markedly between healthy NM-Rs, with higher ranked animals displaying a larger spleen with pro-inflammatory features. NM-Rs with enlarged spleens did not show immature myeloid cells in the peripheral blood as observed in sick NM-Rs with wounds. In all healthy NM-Rs splenic and peripheral blood cell frequency showed an increased myeloid/lymphoid ratio, low bone marrow cellularity and extramedullary hematopoiesis taking place in the spleen with increased erythropoiesis, megakaryopoiesis and myelopoiesis, but reduced B lymphopoiesis compared to mice. B and T lymphocytes were found in secondary sites such as the lymph nodes, gut lymphoid sites and in the thymus, but the latter showed an unexpectedly reduced size in young adults. Our data suggest that, unlike other rodent species, the NM-R spleen is a major site of adult hematopoiesis under normal physiological conditions. However, the reduction in B lymphoid lineage suggests that NM-R immune system relies mainly on innate immune response with a more restricted adaptive immune response.

## Results

### Variable spleen size in NM-Rs

In order to study the structure and function of the NM-R spleen, we collected data from NM-R spleens over the last 4 years from a group of randomly sampled healthy animals (n = 32) aged between 1 and 5 years old, excluding breeding males and queens. Surprisingly, we observed that spleen mass and length varied considerably across healthy NM-Rs (Fig 1A). Spleen size expressed as percentage of body mass (%BM) in C57BL/6N mice (n = 40, aged between 1 and 5 months) was on average 0.32% versus 0.27% in NM-Rs (n = 32). However, spleen size was much more variable in NM-Rs with healthy animals displaying very large or very small spleens (Fig 1A). We divided the NM-Rs into 2 groups based on spleen size frequency distribution that showed dip at around 0.25% of BM (S1A Fig). We classified NM-Rs according to spleen size, NM-Rs with small spleens (ssNM-R: %BM ≤ 0.26%) and large spleens (lsNM-R: %BM > 0.26%) (Fig 1B). The mean spleen mass was 0.18% for ssNM-Rs and 0.35% for lsNM-Rs and the latter showed spleen masses similar to those of mice (Fig 1B). Since the liver and the spleen can both be sites of extramedullary hematopoiesis and could become enlarged during sickness in rodents [17, 18], we also measured liver mass (expressed as %BM) in the same NM-R cohort. Mean liver mass was not different between animals with small and large spleens (Fig 1C). NM-R livers (combined), ssNM-R and lsNM-R were significantly smaller compared to mice (Fig 1C), but the liver size frequency distribution showed a normal distribution in contrast to the spleen size frequency distribution (S1A and S1B Fig). In all NM-Rs the spleen and liver size increased with age, in contrast to the mouse where both organs decreased in size with age (S1C-S1F Fig). The mean age of ssNM-Rs was 28.6 ± 2.3 months versus 36.3 ± 5.7 months for lsNM-Rs which was not significantly different (unpaired t-test: p = 0.06). Spleen size was independent of sex in NM-Rs whereas female mice showed larger spleens compared to males (S1G Fig). Thus, the dynamics of spleen growth in young adult NM-Rs differed considerably from that of age-matched mice. The 32 NM-Rs were taken from 3 distinct colonies (named A, B and C), mean spleen mass was not different between colonies (S1H Fig).

**Fig. 1:**
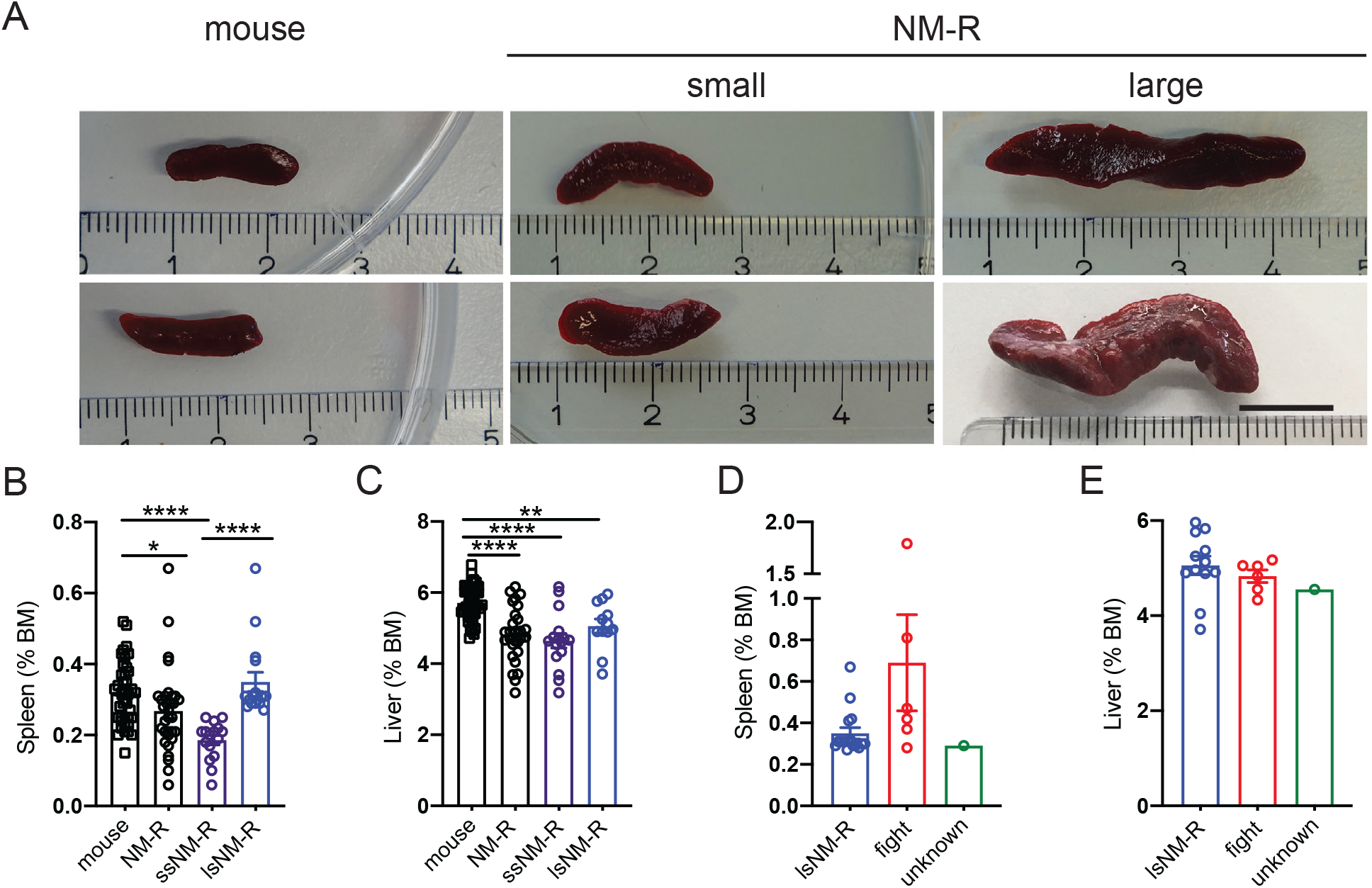
Variable spleen size in NM-Rs. (A) Representative images of NM-R small (ssNM-R) and large (lsNM-R) spleens compared to mouse spleens, scale bar = 1 cm. (B) Spleen weight expressed as % of body mass (% BM) for mice (n= 40) and NM-Rs (n= 32 combined ssNM-R and lsNM-R, n= 16 ssNM-R, n= 16 lsNM-R). (C) Liver weight expressed as % BM for mice (n=40) and NM-Rs (n= 28 combined ssNM-R and lsNM-R, n= 16 ssNM-R, n= 12 lsNM-R). (D-E) Comparison between spleen weight (D) and liver weight (E) of lsNM-Rs (same data as in B-C) and siNM-Rs plotted per type of sickness (fighters n = 6 and unknown cause of sickness n= 1). Percent of BM (% BM) for each tissue type was calculated with BM and tissue weight in g. Graphs represent mean ± s.e.m. Unpaired *t* test: p value *<0.05, **<0.01, and ****<0.0001. See also S1 Fig.

### Splenomegaly in lsNM-Rs is not associated with signs of infection

We next asked whether NM-Rs with enlarged spleens showed signs of ongoing illness or infection, as is the case for other rodents. Enlarged spleen (splenomegaly) may result from extramedullary hematopoiesis in the spleen and liver of individuals suffering from anaemia, neoplasia or myeloid hyperplasia in response to an infection or inflammation. Over a four-year period we collected the spleen and livers from 7 sick NM-Rs (siNM-R). Among them six animals were wounded from fighting and three of these animals had macroscopically infected wounds. These six animals served as positive controls for splenomegaly (cohort named fight or siNM-R fighters). One remaining animal showed signs of sickness, but of unknown cause. The siNM-Rs that had engaged in fights showed the largest spleens (mean = 0.69% of BM) (Fig 1D). The spleen size of siNM-R fighters was twice the size of healthy NM-Rs (combined, mean = 0.27% of BM) or of the lsNM-R (mean = 0.35%) (Fig 1B and 1D), indicating that splenomegaly does occur in NM-Rs following infection. There was no indication of enlarged livers in siNM-Rs regardless of illness type (Fig 1E).

In rodents and humans, increased numbers of immature myeloid progenitors and monocytes in peripheral blood are indicators of infection. Since little is known about the blood cells of NM-R, we first examined bone marrow cells from healthy NM-Rs, the primary site of hematopoiesis in which hematopoietic stem cells generate all immune cells including erythroid, myeloid and lymphoid lineages. NM-R femurs were paler in colour than mouse femurs, suggesting lower haemoglobin and erythrocyte numbers (Fig 2A). Cytospins of bone marrow cells that were not subjected to erythrocyte lysis indicated that all cell types of the erythroid lineage including mature erythrocytes, reticulocytes, orthochromatic erythrocytes and erythroblasts are present in the bone marrow of the NM-R (Fig 2B). In addition, all known hematopoietic cell types found in mouse bone marrow were also present in the NM-R including myeloid and lymphoid lineages (Fig 2B). Surprisingly, in NM-Rs the immature neutrophils (also called band neutrophils) are stab-cell shaped similar to those of humans [21] while characteristic ring-shaped neutrophils of the mouse and rat were not found (Fig 2B). Furthermore, the cell number was 3 times lower in NM-R femur compared to mouse femur (7.3x10^6^ in NM-R versus 25x10^6^ in mouse, Fig 2C). Bone marrow hematopoietic cells differentiate and are found in the peripheral blood from where they can be further characterized. We therefore analysed 23 NM-R blood samples from healthy animals (n = 11 from ssNM-Rs and n = 12 from lsNM-Rs) using an automated blood counter and mouse blood as reference (S2A-S2E Fig). We found similar total white blood cell counts in both species (S2A Fig), but the frequency of the various cell populations was altered with an increased neutrophil/lymphoid ratio and a higher number of monocytes in the peripheral blood of NM-Rs (S2B-S2D Fig). The eosinophil count was only modestly increased (S2E Fig). These blood counts were similar in NM-Rs with small and large spleens, with ssNM-Rs showing slightly lower white blood cell counts (S2A Fig). Thus, our data indicate that the bone marrow of NM-Rs can give rise to all cell types described in mice and humans, but the distribution of the hematopoietic cells in peripheral blood was more similar to that of humans with increased circulating myeloid cells at the expense of circulating lymphocytes.

**Fig. 2:**
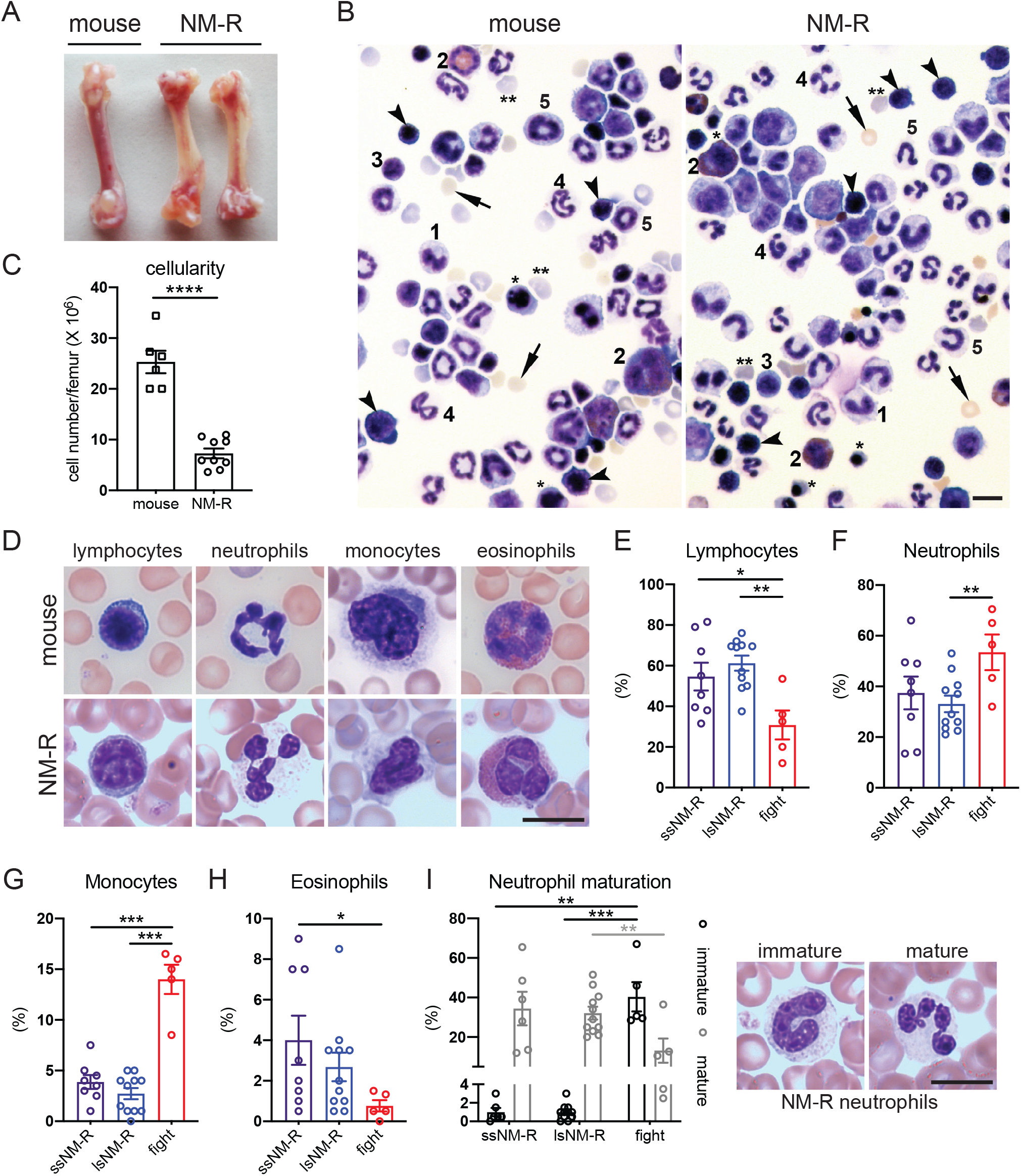
Wounded NM-Rs show high immature neutrophil count in peripheral blood unlike lsNM-Rs. (A) Representative images of femur from mouse and NM-R, (B) May-Grünwald staining of bone marrow cell cytospin without red blood cell lysis. Arrowhead: erythroblast; *orthochromatic erythroblast; **reticulocytes; arrow: mature erythrocytes; 1: monocytes; 2: eosinophils; 3: lymphocytes; 4: neutrophils; 5: band neutrophils. (C) Number of blood cells per femur, n = 6 mice, n = 9 NM-Rs. (D) Representative images of May-Grünwald staining of white blood cells in peripheral blood smears of NM-R and mice: lymphocytes, neutrophils, monocytes and eosinophils. (E-H) Percentage of lymphocytes (E), neutrophils (F), monocytes (G) and eosinophils (H) in the peripheral blood of NM-Rs using May-Grünwald staining of blood smears: n = 8 ssNM-R, n= 11 lsNM-R and n = 5 siNM-R fighters. (I) Left panel: percentage of immature and mature neutrophils counted in blood smear of n = 8 ssNM-R, n= 11 lsNM-R and n = 5 siNM-R fighters. Right panel: representative images of May-Grünwald staining of NM-R immature and mature neutrophils. All scale bars = 10 µm. Graphs represent mean ± s.e.m. Unpaired *t* test: p value *<0.05, **<0.01, ***<0.001 and ****<0.0001. See also S2 Fig.

These data suggested that lsNM-Rs with enlarged spleens are healthy. To verify this, we examined in detail monocytes and immature neutrophil populations in the peripheral blood of animals with infection or inflammation (siNM-R fighters) and compared them to those from lsNM-Rs. Analysis of May-Grünwald stained blood smears from 19 NM-R healthy animals (n = 8 ssNM-Rs and n = 11 lsNM-Rs) confirmed blood counter data (Fig 2D-2H compared to Fig 2E-2H, S2B-S2E Fig). Interestingly, blood smears from siNM-R fighters (n = 5) showed dramatic increases in the monocyte population and a reduction in lymphocytes compared to all NM-R cohorts (Fig 2G and 2E). In addition, 53% of the white blood cells were immature neutrophils (band neutrophils) and 13% were segmented neutrophils (mature stage) in the peripheral blood of the siNM-R fighters, indicating an active immune response against infection or inflammation. In contrast, almost exclusively mature neutrophils (segmented neutrophils: 33 to 41% of white blood cells versus ≤ 1% of immature neutrophils) were found in healthy NM-Rs (ssNM-R and lsNM-R) (Fig 2I). Our results clearly demonstrate that NM-Rs with an apparent splenomegaly do not show myeloid hyperplasia like sick NM-Rs.

### Increased myeloid and reduced lymphoid lineages in NM-R spleen

The differences in spleen size found in healthy NM-R cohorts might reflect specific cell type hyperplasia between both cohorts. To address this, we applied global gene expression profiling to investigate molecular differences between small and large NM-R spleens compared to the mouse. We also compared RNAseq data from mouse spleens with NM-R in order to reveal whether NM-R and mouse spleen share molecular signatures. Global comparison of transcriptomes (including transcripts from 12946 genes) indicated major differences between the two species and high similarity between ssNM-R and lsNM-R (Fig 3A). Principal component analysis and Venn diagram analysis also showed clear species differences and only loose clustering of large and small NM-R spleens (S3A and S3B Fig). Differential gene expression analyses of the 3 groups showed 4869 and 4873 up-regulated genes and 4737 and 4713 down-regulated genes in ssNM-R and lsNM-R, respectively, compared to mice (Fig 3B and 3C). In ssNM-R spleens just 16 genes were differentially up-regulated and 41 were down-regulated compared to lsNM-R spleens (Fig 3B). Comparing NM-R with mouse spleen the RNAseq data revealed a dramatic reduction in the expression of B cell markers (such as *Cd19*, *Cd79a* and *Cd79b*), and of dendritic cell markers such as *Itgax* (coding for CD11c), but there was a dramatic increase in the expression of myeloid markers such as *Itgam* (coding for CD11b) and a modest increase in monocytic/macrophage gene expression (*Cd14*) (S3C and S3D Fig, S1 Table). To infer immune cell type abundance in the spleen from the bulk RNAseq data digital cytometry using CIBERSORT analysis was performed [22]. Since no pre-defined signatures exist for immune cell subsets of NM-Rs, mouse gene expression signatures for the cell subsets were used [22, 23]. The analysis predicted that 53 ± 3% of myeloid cells (31.8 ± 4.6 % granulocytes, 10 ± 1% monocytes and 11 ± 4 % macrophages), 48 ± 3 % of lymphoid cells (10 ± 4% B cells, 2 ± 1 % plasma cells, 7 ± 2% activated NK cells and 29 ± 3 % T cells) are present in NM-R spleens regardless of size (Fig 3D, S3E Fig). This shift towards myeloid cells was highly consistent with recently published single cell RNAseq profiling data from the NM-R [16]. In addition, our functional analysis using gene set enrichment analysis (GSEA) highlighted a significant underrepresentation of gene sets implicated in B cells homeostasis, regulation and proliferation (S2 and S3 Tables). Of note, the percentage of total T-cells was similar in both species, but in NM-Rs the T-cell subset distribution differed from that of mice, including the presence of gamma-delta T-cells and Th1 cells (S3E Fig, left panel). A significant change in the expression of T cell-associated gene sets by GSEA was found (S2 and S4 Tables). Thus, the data suggested that lymphopoiesis and myelopoiesis are differently regulated in adult NM-Rs compared to mice, predicting a special role for the spleen in this species.

**Fig. 3:**
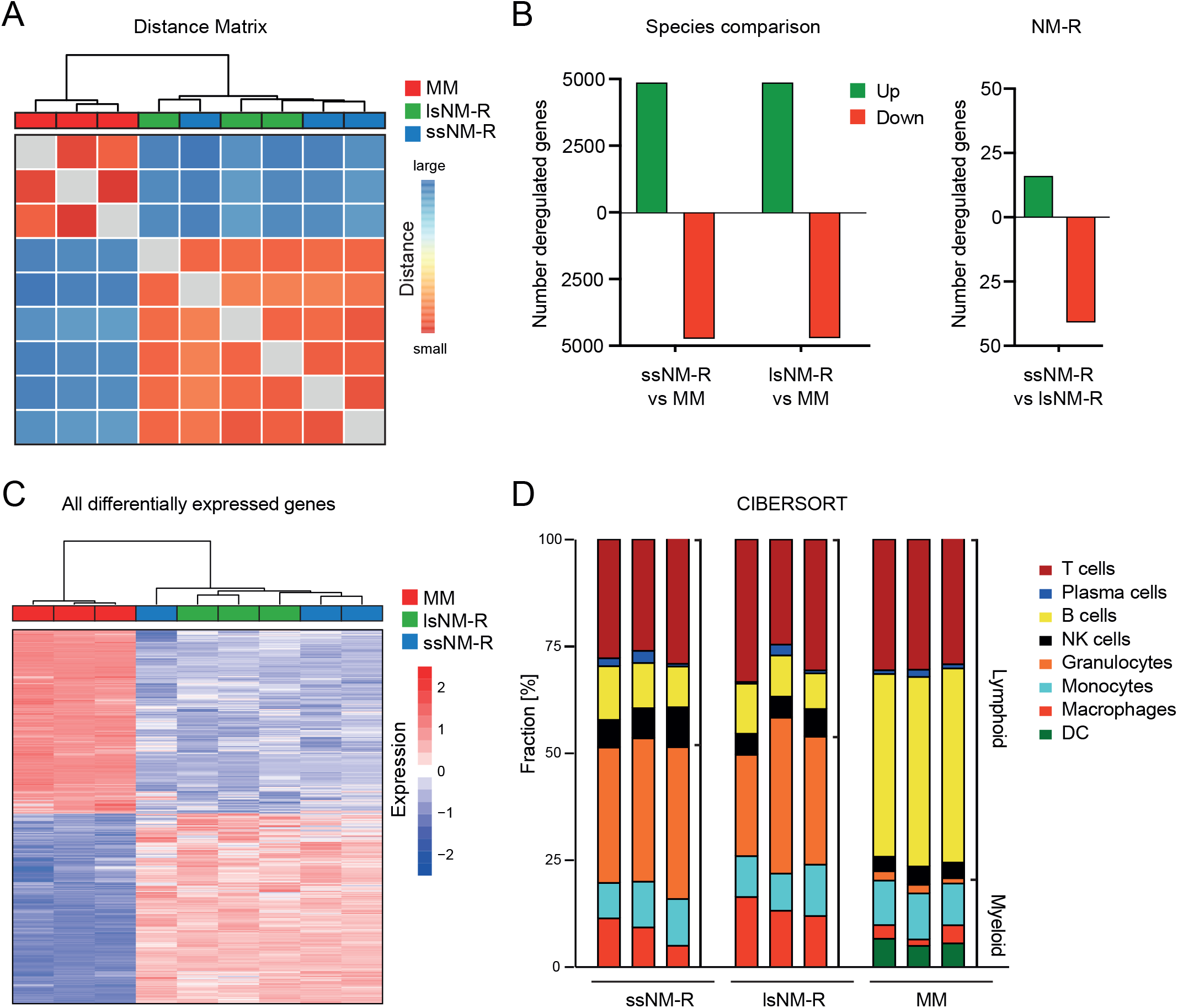
Transcriptomic analysis of NM-R and mouse spleen. (A) Heatmap visualization of the Euclidean distance between spleen samples based on global transcriptomic data (RNAseq). (B) Number of differentially regulated genes (upregulated in green, downregulated in red) in the spleens of ssNM-R, lsNM-R compared to mouse (MM, left panel) or compared to each other (right panel). (C) Heatmap of all differentially expressed genes in the spleen of ssNM-Rs and lsNM-Rs in comparison to mouse. Samples are hierarchically clustered based on Pearson correlation. (D) Fractions of major splenic cell types based on *in silico* cytometric analysis of transcriptomic data (CIBERSORT). ssNM-R: NM-R with small spleen, lsNM-R: NM-R with large spleen, DC: dendritic cells, NK: natural killer cells, MM: mouse (Mus musculus); n = 3 per group. See also S3 Fig.

### The NM-R spleen has unique structural features

The spleen is composed of two functionally and morphologically distinct compartments, the white pulp and the red pulp. The white pulp contains most of the lymphocytes and initiates the immune responses to blood-born antigens while the red pulp is a blood filter that removes foreign material and damaged or senescent erythrocytes, and is a storage site for iron, erythrocytes and platelets [24]. We predicted that, the loss of 30-40% of splenic lymphocytes and the 50% increase in myeloid cells would impact the structure and function of the spleen. Indeed, histological analysis of NM-R spleens showed a strongly reduced white pulp volume and an increase in trabeculae abundance that was independent of spleen size (Fig 4A-4C). We wondered whether these structural peculiarities were observed in other African mole-rat species. We had access to spleens from two other Bathyergidae species the Natal and Highveld mole-rats (*Cryptomys hottentotus natalensis* and *Cryptomys hottentotus pretoriae*) [25, 26] that are related to NM-Rs, but are not eusocial mammals (S4A Fig). The structure of the NM-R spleen did not resemble that of spleens from Natal and Highveld mole-rats, which were both very similar to the mouse and rat (Fig 4A, S4B-S4D Fig). The increased trabeculae density in NM-R spleens was accompanied by a small increase in the expression of the *Col3a1* gene (coding for Type III collagen) a reticulin fibrin component and by a 5-fold increase in the RNA level of α-SMA (encoded by *Acta2* gene) (Fig 4D and 4E). These two genes may be associated with the fibrous trabeculae that act as a pump to filter blood.

**Fig. 4:**
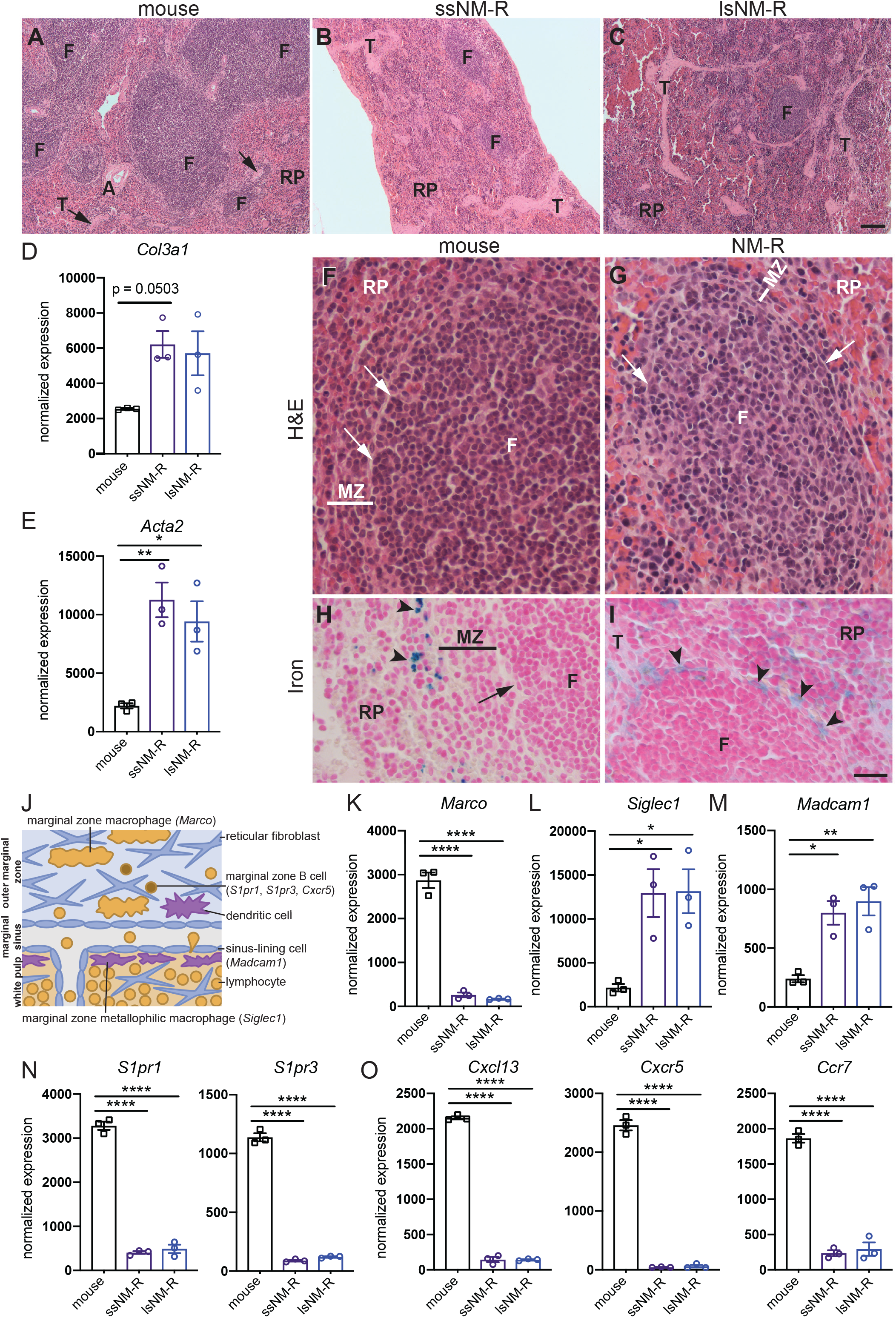
NM-R spleen shows a reduced white pulp compartment and a thin marginal zone. (A-C) H&E staining of the spleen of ssNM-R (B) and lsNM-R (C) in comparison to mouse (A). Note the increase in red pulp/white pulp ratio with reduced number and size of follicles (F) and increased number of trabeculae (T, arrow) in NM-R spleens. (D-E) Increase in normalized RNA expression levels of *Col3a1* (D) and of *Acta2* (E) in NM-R spleens compared to mouse. (F-G) H&E staining of mouse (F) and NM-R (G) spleen showing the presence of follicles (F) with a thinner marginal zone (MZ) in NM-R compared to mouse. (H-I) Iron staining of mouse (H) and NM-R (I) spleen showing stained macrophages (blue iron staining, arrowhead) close to the follicle in NM-R but not in mouse marginal zone. MZ: marginal zone (black or white line), RP: red pulp, marginal sinus (arrow in F-I). (J) Schematic representation of the mouse marginal zone and its cell types with their expression markers. (K-O) Normalized RNA expression levels of (K) *Marco* (MZ macrophage marker), (L) of *Siglec1* (MZ metallophilic macrophage marker), (M) of *Madcam1* (sinus lining cells marker), (N) of markers of MZ B cells (*S1pr1, S1pr3*) and (O) of chemokine-chemokine receptors involved in the lymphocytes migration into the white pulp (*Cxcl13, Cxcr5 and Ccr7*). One-way ANOVA with Tukey’s post-hoc test for multiple comparisons: p value *<0.05, **<0.01 and ****<0.0001. Data is based on RNAseq and bars represent mean ± s.e.m. Scale bar = 100 μm (A-C) and 20 μm (F-I). See also S4 Fig.

The white pulp consists of three sub-compartments: the periarteriolar lymphoid sheath (the T cell zone), the follicles and the marginal zone [24]. In NM-R spleens the follicles were very small and reduced in number compared to mouse spleen (Fig 4A-4C) and were surrounded by a very thin marginal zone (Fig 4F and 4G). The marginal zone is where the blood is filtered from pathogens and is organized in layers with the marginal zone macrophages, the reticular fibroblasts and marginal zone B cells all facing the red pulp. The marginal sinus with its sinus lining endothelial cells and an inner ring of marginal zone metallophilic macrophages separate the marginal zone from the periarteriolar lymphoid sheath and follicles (Fig 4J) [27]. Iron staining labelled red pulp macrophages in mice that are localized to the red pulp (Fig 4H). In NM-Rs iron-stained macrophages were found not only in the red pulp, but also close to the follicles (Fig 4I), suggesting a microarchitectural change of the marginal zone. These anatomical changes were reflected in our RNAseq data that showed decreased *Marco* expression (10-fold compared to mouse), a marker of marginal zone macrophages (Fig 4K), suggesting reduced abundance or loss of marginal zone macrophages. In contrast, there was a 3-fold increase in the expression of marginal zone metallophilic macrophage and sinus lining markers (*Siglec1* and *Madcam1*, respectively) compared to mice (Fig 4L and 4M). The expression of marginal zone B cell receptors (*S1pr1, S1pr3, Cxcr5*) involved in marginal zone B cell migration to the follicles were also decreased probably due to the reduced abundance of marginal zone B-cells (Fig 4N and 4O). Taken together, the low number of marginal zone B cells and marginal zone macrophages could explain the altered morphology of the NM-R marginal zone. This microarchitectural change of the marginal zone might contribute to the impaired adaptive immunity, in particular the proper binding and clearance of blood-borne pathogens. In addition, we found that enlarged spleens of lsNM-Rs did not show signs of pathology associated splenomegaly (Fig 4C).

### Increased splenic granulocytes at the expense of the lymphoid compartment

The formation and maintenance of follicles in lymphoid tissues such as the spleen are regulated by chemokines and their cognate receptors expressed by stromal cells to generate a microenvironment necessary for B and T cell homing to the follicles [28]. The expression levels of chemokine genes (*Cxcl13* and *Ccl19*) and their respective receptors (*Cxcr5 and Ccr7*) were decreased in NM-R spleen compared to mice (Fig 4O, S1 Table). Using the T cell marker CD3e we could show that in NM-Rs T-cells were mainly present in the red pulp, but in much smaller numbers than in mice (Fig 5A and 5B). In both species Western blot analysis showed higher expression of CD3e in thymus (site of T lymphopoiesis) compared to spleen, while no expression was found in the liver, a non-hematopoietic organ (Fig 5E). Furthermore, CD3e expression was lower in NM-R spleens regardless of their size compared to mouse (Fig 5F). Unfortunately, we could not confirm the decrease in B cells in NM-R spleens using immunostaining because of the lack of NM-R specific reagents, but H&E staining rarely showed the presence of follicles and germinal centres, both structures harbouring B cells. These data suggest that splenic adaptive immune responses may rely mainly on T cells in the NM-R.

**Fig. 5:**
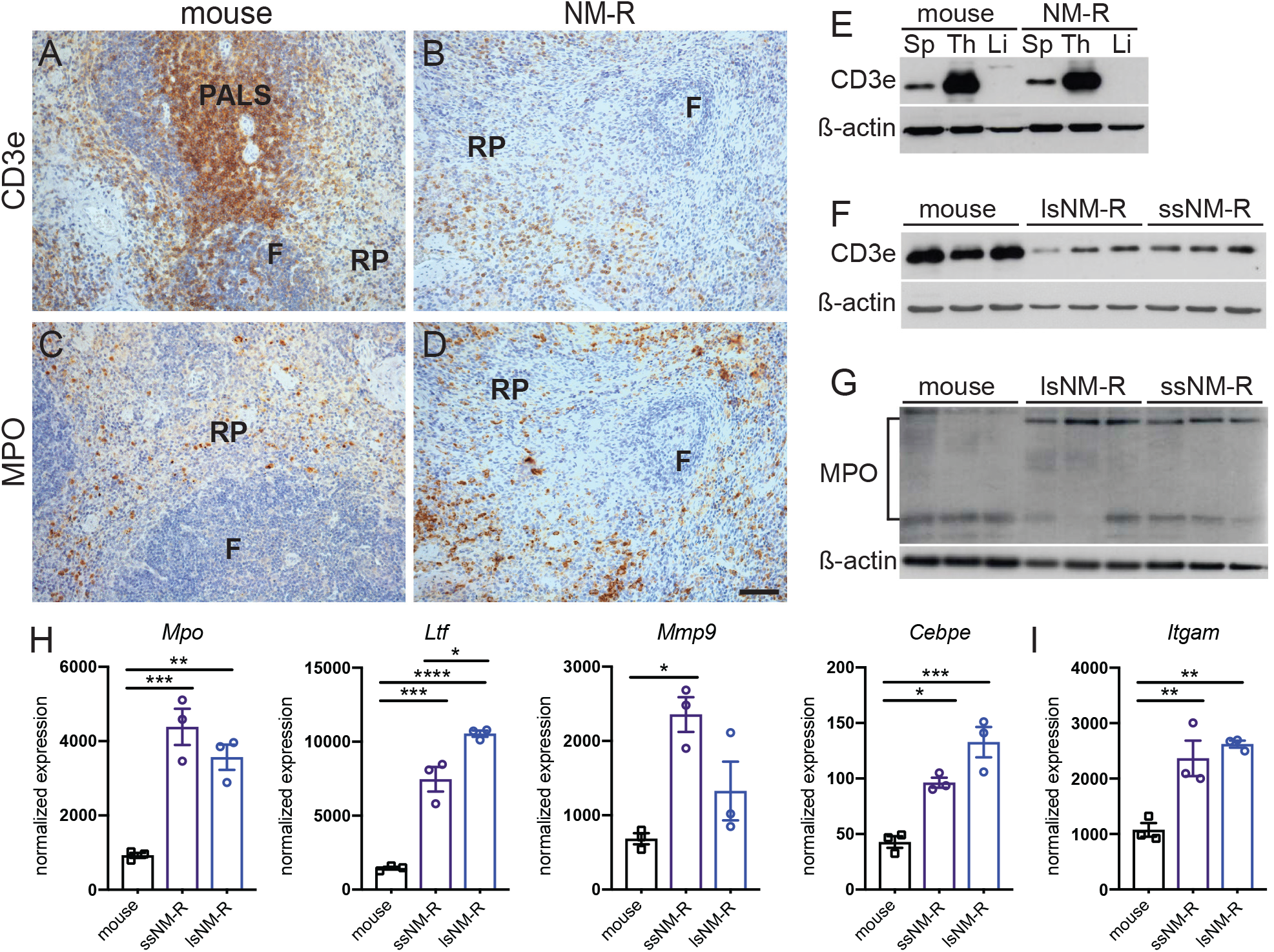
The granulocyte population is increased in NM-R spleen. (A-B) Immunostaining of the splenic T-cells with CD3e antibody and of the splenic granulocytes with myeloperoxidase (MPO) antibody (C-D) in NM-R and mouse. (E) Protein expression of CD3e in various tissues of NM-R and mouse: Sp: spleen, Th: thymus, Li: liver. (F) CD3e (T-cells marker) protein expression in 3 mouse spleens, lsNM-Rs and ssNM-Rs. (G) MPO (granulocyte marker) protein expression in 3 mouse spleens, lsNM-Rs and ssNM-Rs. (H) Increase in normalized RNA expression levels of granulocyte markers (*Mpo, Ltf, Mmp9* and *Cebpe)* and (I) myeloid marker (*Itgam*) in NM-R spleens in comparison to mouse spleens. β-actin expression was used as loading control in E, F and G. F: follicles, PALS: periarteriolar lymphoid sheath, RP: red pulp. Data in H and I are based on RNAseq and bars represent mean ± s.e.m. One-way ANOVA with Tukey’s post-hoc test for multiple comparisons: p value *<0.05, **<0.01, ***<0.001 and ****<0.0001. Scale bars = 50 µm.

Our CIBERSORT and GSEA analyses also predicted increased myeloid populations in particular of granulocytes in NM-R compared to mouse (31.8 ± 4.6% vs 1.8 ± 0.5% total granulocytes including 29.2 ± 4.6% vs 1.4 ± 0.4% neutrophils, and 2.3 ± 1.3% vs 0.4 ± 0.3% eosinophils in NM-R (n=6) vs mouse (n=3), respectively). This was supported by immunohistochemistry and Western blot analyses using an antibody directed against myeloperoxidase (MPO) a marker of pre-and mature granulocytes. We also found higher protein levels of MPO in the splenic red pulp of NM-Rs compared to mice (Fig 5C-5D and 5G). The expression levels of *Mpo* and 3 other markers of granulocytes, *Ltf* (granules of neutrophil granulocytes), *Mmp9* and *Cebpe* (major transcription factor of neutrophil lineage) were increased in NM-R spleens, demonstrating that NM-Rs have more splenic granulocytes than mice independent of size (Fig 5H). Of note, the RNA level of the common marker of myeloid cells, *Itgam* (coding for CD11b) was also increased (Fig 5I). In summary, our data suggest that NM-Rs have enhanced antimicrobial innate immunity, but their adaptive immunity might be less efficient compared to mice.

### Lymphocytes locate to peripheral blood and lymphoid tissues in NM-Rs

Gene expression profiling and histological analysis consistently showed that NM-Rs have almost 50% fewer splenic resident lymphocytes compared to other rodents. This unique immunological feature might have major consequences for adaptive immunity unless compensatory lymphopoiesis occurs in the bone marrow or other secondary lymphoid organs. Lymphocytes were found in bone marrow cytospin and in peripheral blood, but at much lower frequencies than in mice (Fig 2). In contrast to other bone marrow hematopoietic progenitors that undergo several differentiation stages before egression and maturation, T-lymphocyte progenitors migrate to the thymus to differentiate into naïve T-cells that can migrate to the blood and secondary lymphoid organs. Intriguingly, in young adult NM-Rs (1 to 3 years old) the thoracic thymus was often embedded in brown adipose tissue (S5A and S5B bottom Fig) and the thoracic thymus/body mass ratio was considerably lower in NM-Rs compared to young C57BL/6N mice that do not yet show thymus involution (S5B top Fig). Histology of the thoracic thymus revealed a clear cortex and medulla (S5C Fig) in which the naïve T lymphocyte marker CD3e was highly expressed in both mice and NM-Rs (S5D Fig). The NM-R thoracic thymus contains CD3e+ T cells, but its small size prompted us to search for other sites of lymphopoiesis.

Lymphocytes (T and B cells) are also found in lymph nodes. We next analysed NM-R axillary lymph nodes that are 2-4 millimetres in length, similar to those of mice (S6A top panel Fig). Lymph nodes are structurally organized in B and T cell areas, a process regulated by cytokine signalling (S6A, bottom panel Fig). Histologically, B cell and T cell areas were easily identified in axillary and mesenteric lymph nodes of NM-Rs and B cell areas possessed germinal centres (S6B Fig). T cell areas showed high expression of CD3e (S6C Fig).

Since half of the lymphocytes are located in the mucosa-associated lymphoid tissue in mice, we next focused on lymphoid nodules of the small intestine including among others Peyer’s patches [29]. Peyer’s patches are visible to the naked eye in mouse small intestine, but were rarely apparent in NM-R gut (S6D and S6E Fig). In addition, the small intestine of NM-Rs was only a third of the length of that in the mouse, however the length of the colon was similar (S6D and S6F-S6G Fig). Histological analysis of the NM-R small intestine showed lymphoid nodules with a morphology atypical for Peyer’s patches, but these follicles displayed B cell areas with germinal centres containing apoptotic cells and T cell areas expressing CD3e in both NM-Rs and mice (S6H and S6I Fig). In general, we found that NM-Rs and mice have similar lymph nodes with well-structured B and T cell areas, but unlike in mice the NM-R small intestine did not appear to have typical Peyer’s patches, even in sick animals (S6E Fig).

### Increased extramedullary erythropoiesis is likely not a cause of splenomegaly

The unique structural features of the NM-R spleen could not account for the hyperplasic phenotype of the spleen found in the lsNM-R cohort. In mice splenomegaly can be observed under erythropoietic stress such as hypoxia [30, 31]. Since hypoxia is a normal environmental condition for NM-Rs we hypothesized that extramedullary erythropoiesis might occur naturally in NM-R spleen, but more actively in lsNM-Rs spleens. RNAseq analysis and GSEA highlighted a significant enrichment in erythroid gene subsets (S7A and S7B Fig), including markers of early erythroid progenitors such as *Tal1* (erythroid differentiation factor), *Tfrc* (coding for CD71), *Hoxa9* (a marker of pro-erythroblast and basophilic erythroblast) and markers of erythroid precursor proliferation or survival such as *EpoR* (the Epo receptor), *Gata1* and *Bcl2l1* (coding for Bcl-XL) [32–35] (Fig 6A). Thus, early erythroid progenitors appeared more abundant in NM-R spleen compared to mice, regardless of spleen size. Histological analysis revealed erythroid cells in mouse and NM-R spleens often organized in erythroid blood islands (Fig 6B). Immature erythrocytes (enucleated red blood cells also called orthochromatic erythroblasts, see the schematic representation of the erythroid lineage in Fig 6C, top panel) were present in the peripheral blood of NM-Rs (Fig 6C, bottom panel), but not in healthy mice. In addition, mature erythrocytes (RBC) were larger in NM-Rs as indicated by a higher mean corpuscular volume (MCV), but contained less haemoglobin (HGB) and were less abundant in peripheral blood compared to mice (Fig 6D). However, the haematocrit (HCT) was not different between mouse and NM-R (Fig 6D). Together our results showed that extramedullary erythropoiesis occurs in the spleen of both ssNM-Rs and lsNM-Rs, but cannot account for splenomegaly in lsNM-Rs.

**Fig 6:**
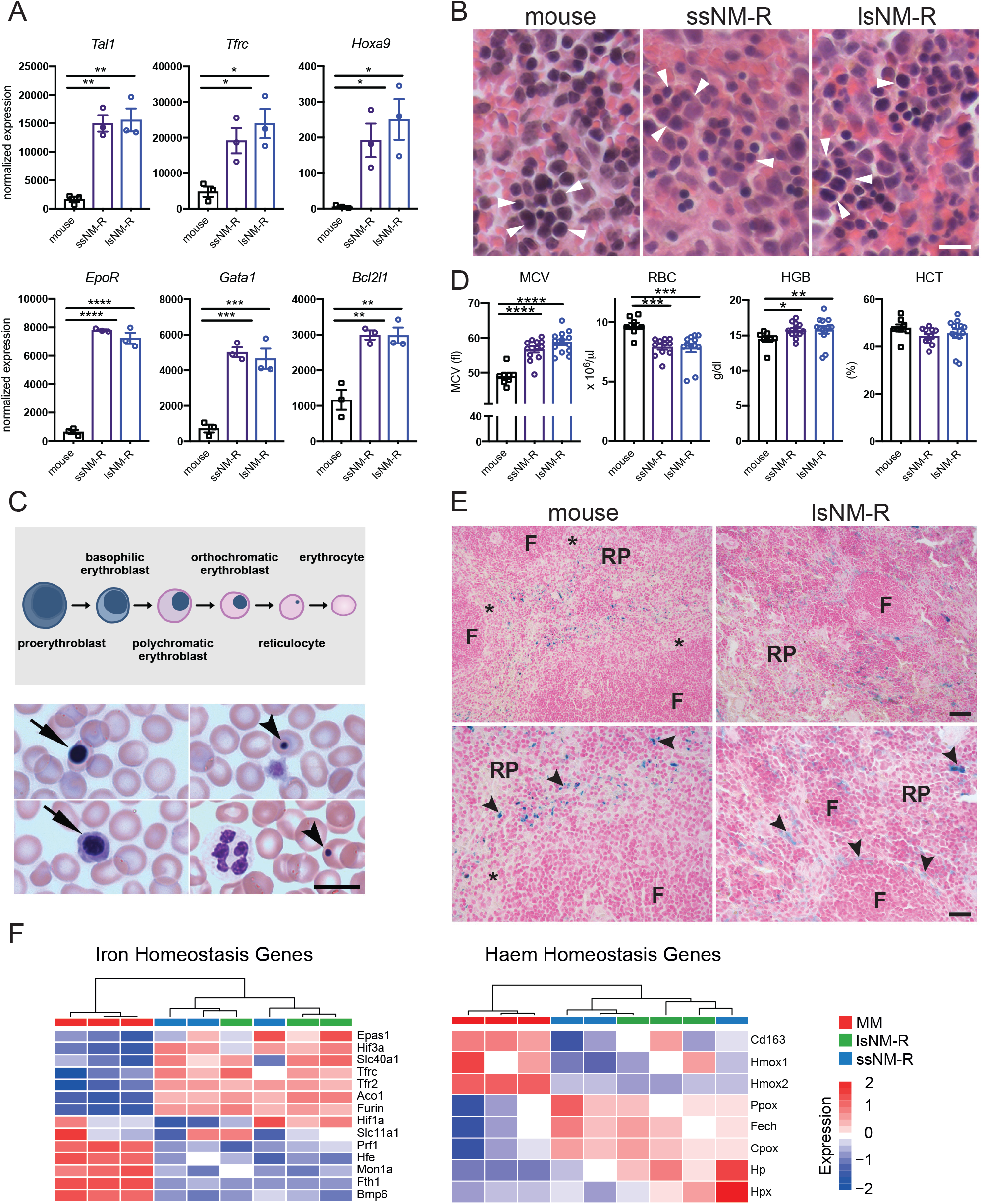
Extramedullary erythropoiesis and iron homeostasis do not explain the hyperplasic spleen of lsNM-Rs. (A) Normalized RNA expression levels of early erythroid progenitors (*Tal1*, *Tfrc*, *Hoxa9*) and of erythroid precursor proliferation and survival (*EpoR*, *Gata1*, *Bcl2l1*) markers in mice and NM-Rs spleens. (B) Representative micrographs of H&E staining of erythroid cells (arrowheads) in the spleen of mouse, ssNM-R and lsNM-R. (C) Schematic representation of late erythroid lineage (top panel). Representative May-Grünwald staining of orthochromatic erythroblast (arrow) in peripheral blood of NM-R, howell-jolly bodies in reticulocytes (arrowhead) in bottom panel. (D) Peripheral blood erythroid parameters in mice (n = 8), ssNM-R (n = 13) and lsNM-R (n = 13). MCV: mean corpuscular volume, RBC: red blood cells, HGB: haemoglobin, HCT: haematocrit. (E) Perl’s Prussian blue staining of ferric iron (arrowhead) in the spleen of NM-Rs and mice. Macrophages close to follicles (F) and in red pulp (RP) contain ferric ion in NM-Rs while in mice only RPM contained ferric ion. Marginal zone: asterisk. (F) Heatmap of genes associated with iron and haem homeostasis in NM-R and mouse spleen. Samples are hierarchically clustered based on Pearson correlation. One-way ANOVA with Tukey’s post-hoc test for multiple comparisons (A) and unpaired *t* test (D): p value *<0.05, **<0.01, ***<0.001 and ****<0.0001. Transcriptomic data is based on RNAseq, MM: mouse (Mus musculus). Data represent mean ± s.e.m, scale bar = 10 μm (B, C), 20 μm (E bottom panels) and 40 μm (E top panels). See also S7 Fig.

### Iron homeostasis is not a cause of splenomegaly in lsNM-R

Thus, NM-Rs probably adapted erythropoiesis to their unusual environmental conditions. The expression levels of several known hypoxia-induced genes (*EpoR, Tfrc, Tfr2, Furin*) were increased compared to the mouse (Fig 6A, S7A Fig, S1 Table), but GSEA analysis did not highlight significant changes in hypoxia-regulated gene subsets (S2 and S3 Tables). In humans and mice erythropoiesis is not only regulated by oxygen availability, but also depends on intracellular iron levels. Indeed, genetic defects in iron or haemoglobin metabolism can lead to splenomegaly in mice [36, 37]. Iron staining was found as expected in red pulp macrophages, cells responsible for efficient phagocytosis of red blood cells and storage of iron [38]. The overall iron accumulation was similar in NM-R spleens independent of their size and compared to mouse (Fig 6E). Intriguingly, we found that the overall GSEA red pup macrophage gene subset was significantly increased in NM-R compared to the mouse (S7C Fig, S4 Table), but RNA levels of *SpiC, Adgre1* (coding for F4/80) and *Cd68*, classical phenotypic markers of mouse red pulp macrophages [39], were strongly decreased in both small and large NM-R spleens (S7D Fig). Lastly our GSEA analyses showed no significant changes in gene subsets implicated in iron and haem homeostasis (Fig 6F, S1-S3 Tables). As a consequence, our data suggest that naturally occurring splenomegaly was not due to defective iron homeostasis.

### Extramedullary megakaryopoiesis does not account for splenomegaly

The erythroid lineage shares a common progenitor with megakaryocytes, the so-called megakaryocyte-erythroid progenitors (MEP) which give rise to erythroid and megakaryocyte lineages in bone marrow (S8A Fig). We hypothesized that extramedullary megakaryopoiesis might also occur in the NM-R spleen. RNAseq analysis indicated that expression of many megakaryocyte and platelet genes are upregulated in NM-Rs compared to mice independent of spleen size (S8B Fig). Megakaryocyte differentiation occurs in several steps from MEP to activated megakaryocytes that release platelets into the peripheral blood [40] (S8A Fig). The RNA levels of marker genes of MEP and early megakaryocytes (*Cd34*, *Gfi1b*, *Fli1*, *Itga2b*), terminal differentiation genes (*Nfe2*, *Tubb1*) and platelets (*Gp1bb*, *Itgb3*, *Cd63*) were all elevated in NM-R spleens compared to that of mice (S8B Fig). Histological analysis also showed a significant increase in the number of megakaryocytes per unit area in lsNM-Rs compared to the rat spleen and a slight increase in megakaryocyte number in NM-R (combined) compared to those of the mouse and rat (S8C and S8D Fig). Surprisingly, blood counts indicated that platelets were less abundant in the peripheral blood of all NM-R cohorts compared to mice (S8E Fig), but NM-R platelet size was larger as shown by mean platelet volume (S8F Fig). Blood smear analysis showed the presence of immature and mature platelets in NM-R peripheral blood, whereas only mature platelets were observed in mice (S8G Fig). Intriguingly, the expression levels of regulatory genes of megakaryocyte terminal differentiation (*Ccl5*, *Il1a* and *Igf1*) were reduced compared to the mouse (S8H Fig). Taken together our data suggest that megakaryocytic differentiation occurs efficiently in NM-Rs, but the terminal differentiation step(s) might be regulated differently. The presence of immature platelets in peripheral blood of NM-Rs might reflect a thrombocytopenia-like phenotype as observed in thrombocytopenia or inherited diseases in humans [41].

### Hyperplasic spleens are associated with higher rank

We detected a small set of differentially expressed genes between NM-Rs with large and small spleens (41 upregulated genes and 16 downregulated) (Fig 3B, S9 Fig). Prediction of immune cell distribution using CIBERSORT analysis showed a specific difference in the apparent incidence of the M0 macrophage subtype (2.8 ± 1.5% in ssNM-R versus 10.4 ± 3.4 % in lsNM-R) (Fig 7A). Furthermore, GSEA highlighted significant increases in hallmark gene subsets such as inflammatory response, granulocytes, naïve T- and B-cells pathways and the Ltf-high-neutrophil subset in lsNM-R compared to ssNM-R (Fig 7B and 7C). These findings suggested that NM-Rs with larger spleens are better equipped for defence against pathogens. Indeed, larger spleen size might confer a survival advantage for NM-Rs. NM-Rs are eusocial mammals with a structural hierarchy with the queen and her consorts occupying the highest rank [12]. We modified a ranking index of animals in a colony with the highest ranking set to 1 for the queen and the lowest to 0 for subordinate [13, 42]. The ranking index of 20 healthy NM-Rs (ssNM-R, lsNM-R) used in this study had been determined (Fig 7D). Strikingly most of the animals (62.5%) with a small spleen were found to have the lowest rank (Fig. 7D). In contrast, many more of the animals with large spleens belonged to the higher ranks (Fig 7D). We found a significant positive correlation between ranking index and spleen mass (%BM) when combining all healthy cohorts (ssNM-R, lsNM-R) (Fig 7E). In contrast, the liver mass (%BM) was poorly correlated with the ranking index in both cohorts (Fig 7F). There was a significant correlation between rank and BM (Fig 7G), but the age of the animals was a poor predictor of ranking or BM (Fig 7H and 7I). Our data identify rank as being predictive of spleen size in healthy animals with large spleens conferring an immunological advantage over lower ranked animals.

**Fig 7:**
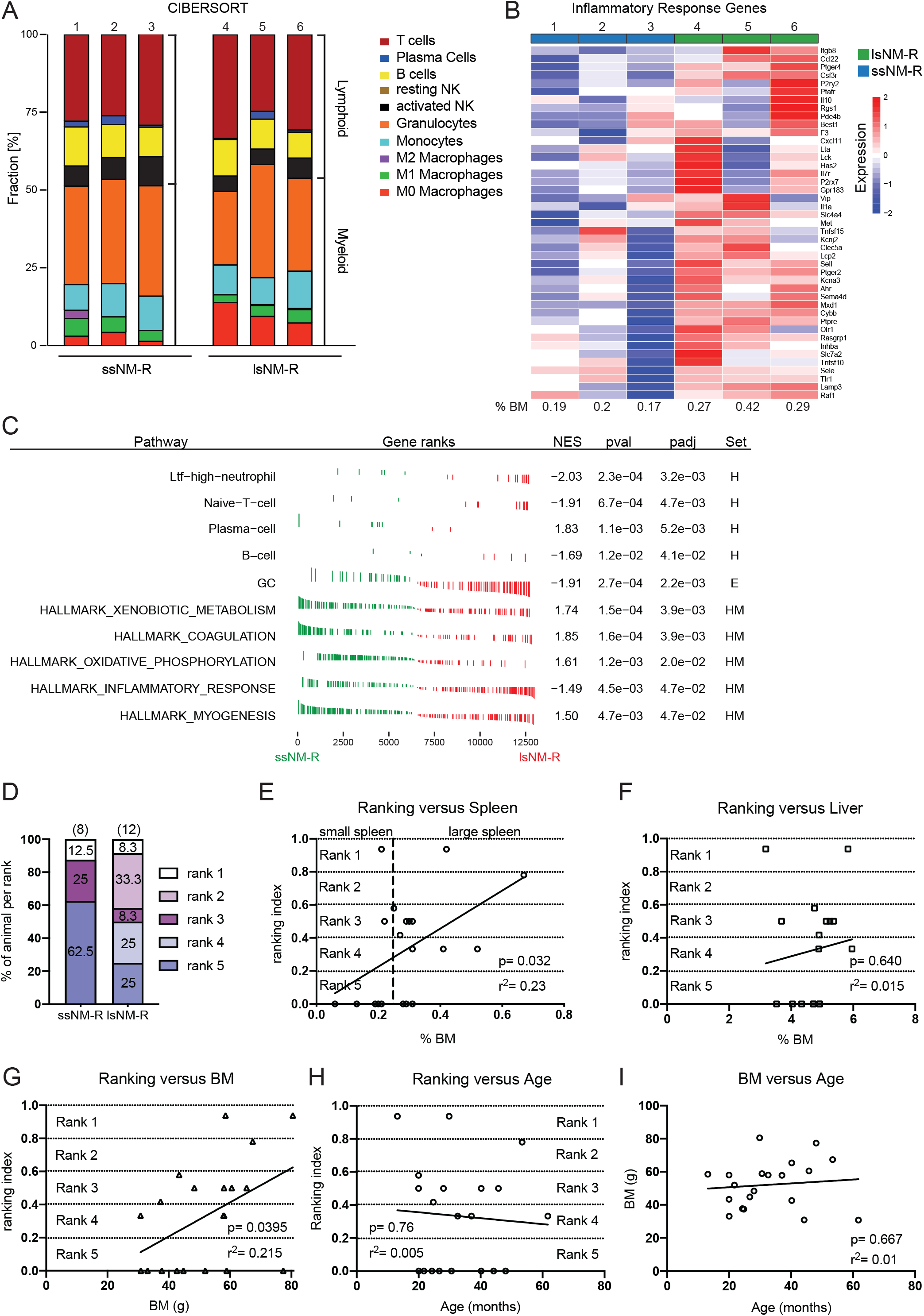
Hyperplasic spleens in lsNM-R are associated with higher rank. (A) Relative fraction of immune cells predicted by CIBERSORT in ssNM-R and lsNM-R spleens (n = 3 per cohort). NK: natural killer cell. (B) Heatmap representation of leading edge inflammatory response genes for ssNM-R and lsNM-R spleens identified by hallmark GSEA analysis. (C) GSEA of transcriptomic data from ssNM-R and lsNM-R spleens (n = 3 per cohort). NES, normalized enrichment score; pval, p value; padj, adjusted p value. Significantly enriched pathways using NM-R cell type signatures from Hilton and colleagues (“H”) and Emmrich and colleagues (“E”) [16, 43] and established hallmark pathways (“HM”). GC: granulocytes. (D) Percentage of animal per rank and per cohort. Note that only 25 % of the lsNM-R has the lowest ranking index (rank 5) compared to more than 60% in ssNM-R. (E-F) Correlation between rank of NM-R in their colony (ranking index) and either spleen size (%BM) (E), liver size (%BM) (F), body mass (BM) (G) or age (H) showing that the ranking index of NM-Rs positively correlated with their spleen size and BM but not with their liver size and age. Body mass and age of NM-Rs poorly correlated (I). Simple linear regression analysis was used to test goodness of fit: the calculated r square (r^2^) and p value are given for each line. NM-Rs n = 20 including n = 8 ssNM-R, n = 12 lsNM-R. See also S9 Fig.

## Discussion

An enlarged spleen may indicate that the immune system is reacting to infection in rodents [17, 18]. In our survey of the immune system of NM-Rs we found an unusual variation in spleen size in apparently healthy individuals. We found that healthy animals with enlarged spleens showed similar white blood cell composition and splenic structural features to animals with small spleens. In contrast, NM-Rs suffering from wound infection displayed enlarged spleens with accompanying signs of immune activation like increased blood monocytes and increased numbers of immature neutrophils (Fig 2G and 2I). Interestingly, the spleens of sick animals were on average larger than those of lsNM-Rs, but this was not statistically different (Fig 1D). We could show that in healthy animals both small and enlarged spleens were associated with enhanced erythropoiesis, megakaryopoiesis, and with myeloid hyperplasia when compared to mice (Fig 5-6, S8 Fig). However, we detected significant molecular differences between small and enlarged spleens of healthy NM-Rs, with large spleens harbouring larger numbers of an LPS-responsive granulocyte population (also called Ltf-high-neutrophil), recently described in NM-Rs [16]. Intriguingly, the molecular profile of larger spleens suggested a pre-activated state that might prepare the animal better for fighting infection. One unusual feature of NM-R colonies is that they display a hierarchical structure with the highest ranked members most likely to be or become breeders [12]. Indeed, it has also been shown that higher ranked NM-Rs such as breeders show longer lifespans compared to lower ranked individuals [4] as well as better survival rates following viral infection compared to non-breeders [14] (V. B unpublished data). Furthermore, higher-ranked individuals are often tasked with colony defence, hence these individuals have a higher risk of coming into contact with intruders which may carry pathogens [12]. Body mass is positively correlated with rank in NM-Rs [13, 42], and here we extend this finding by showing that spleen size also positively correlates with the rank of animals. Interestingly, the size of other organs like the liver showed no correlation with the animal’s rank. Our data suggest that when an animal attains a high rank within the colony this is associated with a change in spleen size and its molecular make-up. We propose that NM-Rs with enlarged spleens may have a survival advantage over lower ranked animals. The immunological repertoire of animals with large spleens may help them to better fight infection, or could even confer cancer resistance. Thus, we have shown a remarkable plasticity in the immune system of NM-Rs that may be regulated through social interaction. When members of the colony get sick social distancing as practised by some species [44] may not be feasible. Thus, tuning of immune competence in higher ranked NM-Rs may be a novel strategy in the animal kingdom to deal with the challenge of infection in a tightly knit colony. The factors that drive spleen plasticity remain to be determined.

We also compared the anatomical and molecular properties of the NM-R spleen to those of other rodents including mice, rats and two other social members of the Bathyergidae family to which NM-Rs belong [26]. Our analysis showed that the NM-R spleen has several unique features compared to other rodents. In agreement with recent studies we showed a dramatic reduction in B lymphoid lineage in the spleen resulting in a decreased white pulp/red pulp ratio [16, 45]. Splenic B lymphocyte content was low in all NM-Rs used in this study, including sick NM-Rs. Furthermore, we found that NM-R spleens exhibit a unique microarchitecture of the marginal zone (reduced marginal zone B cells and marginal zone macrophage populations) which might indicate that the clearance of blood-borne pathogens may be altered compared to other species. This observation is consistent with data from the literature indicating that NM-Rs readily succumb to herpes or coronavirus virus infections [14, 15]. We also show that lymphopoiesis in NM-Rs is maintained since lymphocytes were found in peripheral blood, lymph nodes, mucosa-associated lymphoid tissues in the gut and the thymus. The thoracic thymus of NM-Rs was much smaller than those of mice at different ages. However, CD3+ T cells were found in all lymphoid organs, suggesting that T cell lymphopoiesis takes place in the thoracic thymus. Cervical thymus has been described in NM-Rs and other mammals, including mice and marsupials [15, 46], where T cell lymphopoeisis can occur, however we did not analyse cervical thymus in the NM-Rs examined here. It is well known that intrinsic and extrinsic factors such as cytokines and chemokines are involved in T and B cell migration and homing [28, 47]. Our present data show that RNA levels of such factors (*S1pr1, S1pr3, Cxcl13, Cxcr5, Ccr7, Lta, Nkx2-3* and *Ctsb*) were reduced in NM-R spleens (S1 Table). The mechanisms of tissue homing specificity observed in NM-R and the factors involved in this process remain to be determined.

We did not find reports describing viral or bacterial infections of NM-Rs in the wild [48–51], unlike their close relatives of the genus *Cryptomys* and *Bathyergus* that harbour Bartonella [52]. However, in captivity NM-Rs have been reported to be susceptible to coronavirus infection [14] and in our own laboratory we lost more than 55% of 2 colony members (in colony 1: 20 out of 35 NM-Rs and in colony 2: 5 out of 8 NM-Rs) within a few months because of an unknown viral infection (data not shown). Interestingly, in both laboratories after these mass die-off events almost all queens survived. The relative susceptibility of NM-Rs to viral infection may be due to a narrower immune cell spectrum available to eliminate pathogens with reductions in B cell lineages, dendritic cells, marginal zone macrophages and canonical NK [16] (and our present work). This is in contrast to observations of viral tolerance in some long-lived bat species [53]. However, we found among NM-R splenic immune cell repertoire interesting cells such as gamma-Delta T cells, a special T lymphocyte subset known to be at the border between evolutionary primitive innate system and the adaptive immune system (S3E Fig). Gamma-delta T cells play a role in the “first line of immune defence” against viruses, bacteria and fungi [54, 55]. The presence of more neutrophils and an LPS-responsive granulocyte population also support the idea of enhanced anti-bacterial defences in NM-Rs. Intriguingly, in sick NM-Rs despite increased numbers of immature neutrophils in peripheral blood indicating emergency myelopoiesis in response to injury [56], the animals did not recover, some even developed abscesses, suggesting increased vulnerability to secondary infection probably partially due to lymphopenia.

We also show that adult hematopoiesis takes place in NM-R spleen in addition to adult bone marrow hematopoiesis under normal physiological conditions, and regardless of spleen size. In rodents extramedullary erythropoiesis is observed in response to hypoxia [30, 57]. Thus, the active hematopoiesis in the spleen might reflect an adaptation of the NM-R to compensate for hypoxic environments. Surprisingly, despite the increase in splenic megakaryopoiesis a thrombocytopenia-like phenotype is observed in the peripheral blood of NM-Rs with low platelet counts and the presence of immature platelets. This could also be due to the hypoxic habitat of NM-Rs since in mice hypoxia induces thrombocytopenia [58].

Interestingly, the NM-R immune system displays more similarities to humans than to that of other rodents with a larger myeloid compartment in peripheral blood and spleen and insignificant splenic lymphopoiesis with gamma-delta T-cells. Indeed, NM-R immature (stab-shaped neutrophils) and mature neutrophils found in bone marrow and in peripheral blood resemble those of humans [59]. Food, body size and physiology are factors known to influence spleen development [60]. We also found that thoracic thymus development was quite distinct in the NM-R compared to mouse. Hormonal and endocrine status can influence the development of the immune system [61] and it should be noted in this context that all NM-Rs used in this study were non-breeders and therefore reproductively suppressed [42]. We, like others have found low RNA levels of NK markers (*Ncr1, Nkg7* and *Gzma*). Furthermore, *Adgre1* expression (coding for F4/80), a known rapidly-evolving gene in monocytic/macrophage lineages [62] and a marker of liver resident-macrophages and red pulp macrophages in mice, was almost absent in the spleen and liver of NM-Rs (S7D Fig and data not shown). This suggested that evolutionary pressure selected against the expression of such genes in the NM-R. Indeed, differences in phenotypic marker expression of immune cells between NM-R and mice should be treated with caution. Bone marrow macrophages of NM-R express the NK1.1 receptor of NK cells and are activated by NK1-1 antibodies *in vitro* [63]. We found low RNA levels of mouse classical red pulp macrophages markers (F4/80, *SpiC*, *Cd68*) (S7D Fig), however, GSEA and histological analyses showed that macrophages are present in the red pulp and they store iron (Fig 4E and 6I, S4 Table). Whether these macrophages resemble mouse red pulp macrophages remains to be determined. Interestingly, development and survival factors characteristic of the murine red pulp macrophages (*Irf8, Irf4, Bach1*) [39] were inversely expressed in NM-Rs compared to mice (S7D Fig, S1 Table). Unfortunately, we could not validate our data obtained on B cells, dendrite cells and macrophages due to a lack of specific reagents recognizing these immune cells in the NM-R.

Age-related changes of the immune system in humans and mice are thought to be caused by reduced thymus activity and chronic low-grade inflammation caused by increased activity of the innate immune system [64]. Interestingly, the composition of the NM-R spleen in healthy young animals is reminiscent of that of aged mice, including reduced abundance marginal zone macrophages [65]. It remains to be seen whether the NM-R immune system is better equipped to prevent oncogenic events. Our observations of molecular and anatomical plasticity of the spleen in healthy higher ranked animals raise the intriguing possibility that social success in this species may recruit the immune system to promote longevity.

## Materials and methods

### Animals

Healthy naked mole-rats (aged between 1 and 5 years, e.g., adolescent and young adults) and sick NM-Rs (aged between 1,7 and 7 years) were housed at the Max-Delbrück Center (MDC) in Berlin, Germany, in cages connected by tunnels, which were contained within a humidified incubator (50-60% humidity, 28-30°C), and heated cables ran under at least one cage per colony to allow for behavioral thermoregulation. Food (sweet potato, banana, apple, and carrot) was available ad libitum [66]. NM-Rs were sacrificed by decapitation.

Adult non-reproductive Natal mole-rats (*Cryptomys hottentotus natalensis*) and Highveld mole-rats (*Cryptomys hottentotus pretoriae*) were housed at the Department of Zoology and Entomology, University of Pretoria, South Africa in temperature-controlled rooms set at 25°C and a photoperiod of 12L:12D. The humidity in the rooms was around 40-50%. Mole-rats were fed on chopped vegetables and fruit daily and cleaned weekly with fresh wood shavings and paper towelling. All animals were humanly euthanized by decapitation under EC014-17.

Mice (aged between 4 weeks to 5 months, e.g., young and adults) were fed *ad libitum* with standard diet and water on a 12-hr light-dark cycle at 22°C ± 2°C under 55% ± 10% humidity. Mice were housed in a pathogen-free facility at the MDC, Berlin, Germany. All procedures and animals experiments were conducted in compliance with protocols approved by the institutional Animal Care and Use Committee Landesamt für Gesundheit und Soziales Berlin (LAGeSo). Mice were sacrificed by cervical dislocation. All efforts were made to minimize animal suffering.

### Blood count, blood smear and bone marrow cell cytospin staining

Blood was collected after decapitation of NM-Rs directly into EDTA-containing tubes. Blood from mice was drawn via cardiac puncture and immediately transferred into EDTA-containing tubes. Blood cell counts were measured with an automated veterinary hematological counter Scil Vet abc (SCIL GmbH, Viernheim, Germany) or IDEXX ProCyte Dx hematology analyzer (IDEXX, Germany) with software optimized for mouse blood parameters. May-Grünwald staining of blood smears was performed according to the manufacturer protocol (Sigma, Germany) and the cell type counts of the white blood cells were determined using a Leica DM 5000 B with a x100 oil objective. At least 200 white blood cells were analysed per animal.

For performing cytospin and determining femur cellularity, bone marrow cells were flushed out from the femur, mechanically dissociated and counted using a TC20 automated cell counter (BioRad). For cytospin, 100.000 cells were centrifuged onto slides using a centrifuge slide stainer (Wescor) and stained manually with May-Grünwald staining.

### H&E, Iron staining and immunostaining

Spleen, thymus, lymph nodes and small intestine Swiss rolls were rapidly collected, fixed overnight in 4% paraformaldehyde, embedded in paraffin, sectioned at 4 μ and stained with hematoxylin & eosin histological stain according to the standard protocol. The histological detection of ferric iron in the spleen was performed using an iron staining kit (ab 150674, Abcam).

For immunostaining, sections were deparaffinized and submitted to antigen retrieval (Citrate buffer pH 6) using a microwave. After 2 washes with TBS-T (TBS with 0.05% Tween 20), sections were blocked with TBS-T + 5% goat serum for 30 min at room temperature, and then incubated with rabbit primary antibodies overnight at 4°C. Primary antibodies were diluted in TBS-T + 1% goat serum. Sections were then washed 3 times with TBS-T, subsequently incubated with goat anti-rabbit-HRP (111-035-003, Jackson Immuno-research) for 1 h at room temperature. The rabbit primary antibodies CD3e (1:200, ab 5690, Abcam) and MPO (1:500, A0398, Dako) were used. Dako-EnVision+System-HRP (K4002, Dako) was used for immunodetection. Hematoxylin counter staining was performed before mounting. All images were acquired using a Leica DM 5000 B. To quantify the number of splenic megakaryocytes, four randomly chosen fields in red pulp were photographed at 40X magnification for each animal and analyzed using Item 5 software program (Version 5).

### RNA preparation and RNA sequencing

Total RNA was isolated from three biological replicates per species and per group using RNeasy extraction kit (Qiagen). RNA-seq libraries were prepared using the Truseq Stranded total RNA kit (Illumina) and sequenced on the Illumina HiSeq2500/4000 platform according to the manufacturer’s instruction at Macrogen (Macrogen, Korea). Reads were aligned to mm9 and hetGla2/hetGla Female_1.0, respectively, using STAR aligner version 2.5.3a. The aligned reads were then transformed to raw count tables using htseq-count version 0.10.0. The raw and normalized data are deposited at Gene Expression Omnibus (GEO, accession number GSE179350).

### Transcriptomic analysis

Pre-processed RNA-seq data were imported in R (v3.5.1) for downstream analysis. NM-R genes were annotated to *Mus musculus* homolog associated gene names using the biomaRt package (v2.38.0) to allow merging of the mouse and NM-R data sets. Genes were pre-filtered to remove those transcripts not corresponding to gene symbols or not reaching read sums higher than 10 across all samples. The DESeq2 package (v1.22.2) served for normalization and differential expression analysis. Differentially expressed genes were called using a threshold of an adjusted p value (padj) < 0.05 after multiple-testing correction (Benjamini-Hochberg). For global expression analysis principal component analysis was done using pcaExplorer (v2.8.1) based on the top 3000 variable genes and a global distance matrix was generated using the Euclidean distance. Expression of gene sets was visualized using the pheatmap package (v1.0.12), gene wise scaling and Pearson correlation as distance measure for hierarchical clustering where applicable. To perform gene set enrichment analysis the fgsea package (v.1.8) was used and pre-built, established gene sets were applied (http://www.go2msig.org/cgi-bin/prebuilt.cgi) or custom gene sets were generated from published data derived from NM-R transcriptomes [16, 43]. To assess the cellular composition within the spleens based on the bulk transcripome data, CIBERSORT analysis was performed using the web interface as described by Newman and colleagues [22]. For this, mouse immune gene expression signatures were used as presented by Chen and colleagues [23].

### Immunoblotting

Tissues were lysed with 8 M urea and protein analyzed by SDS/PAGE/protein blotting using rabbit antibody against CD3e (ab 5690, Abcam), rabbit antibody against anti-human MPO (A0398, Dako), mouse β actin (A1978, Sigma), and horseradish peroxidase-conjugated secondary antibodies (111-035-003, Jackson Immuno-research), and chemiluminescence detection (Thermo Fischer).

### Hierarchy assessment and ranking index

Methods as described in [13] and modified from [42]. In brief, two NM-Rs were allowed to approach each other head on in an artificial plastic tunnel. During these interactions the more dominant individual will reliably climb over the subordinate individual. Using a single elimination strategy, with a minimum of three trials for each pseudo-randomly selected pairing of NM-Rs from a single colony, a ranking index (R.I.) was calculated for each colony. R.I = (number of wins) divided by (the total number of behavioural trials). R.I. values were normalized to the maximum value for each colony and the following rankings were assigned based on R.I.: rank 1, R.I. > 0.8, rank 2, R.I. > 0.6, rank 3, R.I. > 0.4, rank 4, R.I. > 0.2, rank 5, R.I. < 0.2. The queen was assigned a rank of 1.

### Statistical analysis

All data are expressed as mean ± s.e.m. Data were first tested for normal distribution. For CIBERSORT analysis variation is reported as ± standard deviation. Statistical tests performed can be found in the figure legends. Statistical analyses were carried out using (Prism 8, GraphPad Software) unless otherwise stated. P value<0.05 was considered to be statistically significant.

## Acknowledgements

We would like to thank U. Höpken (MDC) for her comments on the manuscript. We thank K. Zimmermann (MDC) for converting raw sequencing data, M. Strehle for the graphics, M. Braunschweig and F. Bartelt for excellent technical assistance, P. Langner, S. Schelenk and A. Heuser for automated blood count measurement (PRC, MDC), and G. Pflanz, A. Mühlenberg and I. Duckert for excellent care of the naked mole-rats.

## Author contributions

Conceptualization: Valérie Bégay, Gary R Lewin

Data curation: Valérie Bégay, Branko Cirovic

Formal analysis: Valérie Bégay, Branko Cirovic

Funding acquisition: Gary R Lewin

Investigation: Valérie Bégay, Branko Cirovic, Alison J Barker, Gary R Lewin Daniel W Hart, Nigel C Bennett

Methodology: Valérie Bégay, Branko Cirovic, Alison J Barker

Project administration: Valérie Bégay

Resources: Gary R Lewin

Supervision: Valérie Bégay

Validation: Valérie Bégay, Branko Cirovic, Robert Klopfleisch

Visualization: Valérie Bégay, Branko Cirovic

Writing-original draft: Valérie Bégay

Writing-review & editing: Valérie Bégay, Branko Cirovic, Alison J Barker, Daniel Hart, Nigel C Bennett, Gary R Lewin

## Abbreviations

BM: body mass
GSEA: gene set enrichment analysis
H&E: hematoxylin and eosin
MEP: megakaryocyte-erythroid progenitors
MPO: myeloperoxidase
NK cells: natural killer cells
NM-R: naked mole-rat
RNAseq: RNA sequencing
ssNM-R: naked mole-rat with small spleen
lsNM-R: naked mole-rat with enlarged spleen
siNM-R: sick naked mole-rat

## Competing interests

The authors have declared that no competing interests exist.

**S1 Table: Differential gene expression analysis of NM-R and MM spleen transcriptomes**

DESeq2 output tables for differential gene expression analysis comparing spleen samples from lsNM-R versus MM (“lsNM-R_vs_MM”), ssNM-R versus MM (“ssNM-R_vs_MM”) and ssNM-R versus lsNM-R (“ssNM-R_vs_lsNM-R”), respectively. baseMean, average of the normalised count values; lfcSE, standard error estimate for log2FoldChange; stat, Wald statistic; padj, adjusted p value. For more details, see methods section.

**S2 Table: Gene set enrichment analysis**

GSEA results comparing ssNM-R versus MM spleen transcriptomic data sets. padj, adjusted p value; ES, enrichment score; NES, normalised enrichment score; nMoreExtreme, number of more significant random gene pathways. Pathway size: number of tested genes in the pathway. For more details, see methods section.

**S3 Table: Hallmark gene set enrichment analysis**

GSEA results comparing ssNM-R versus MM spleen transcriptomic data sets focusing on hallmark pathways. padj, adjusted p value; ES, enrichment score; NES, normalised enrichment score; nMoreExtreme, number of more significant random gene pathways. For more details, see methods section.

---

**Fig S1:**
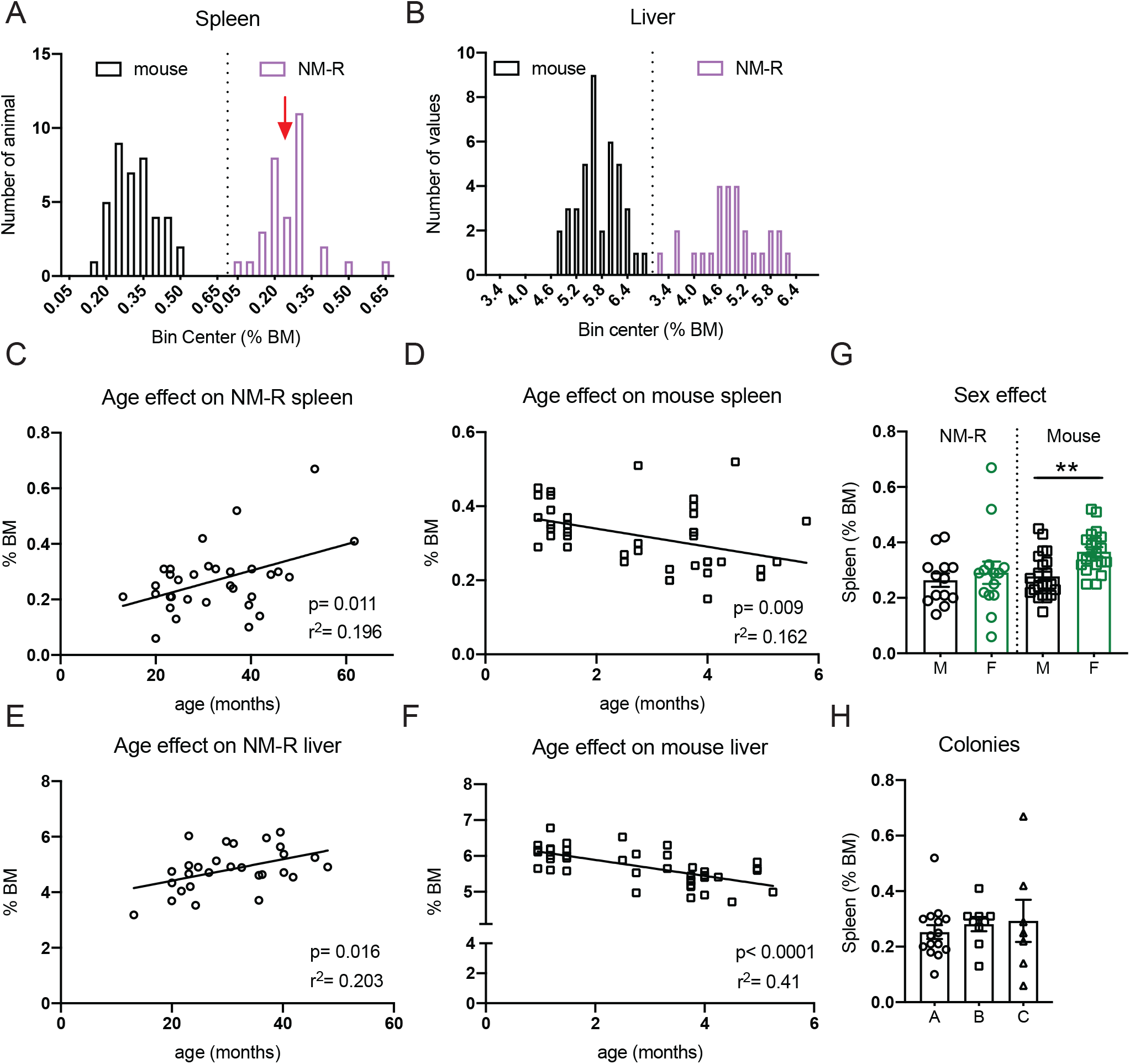
Related to Fig.1. Variable spleen size in NM-Rs. (A-B) Frequency distribution of spleen mass (A) and liver mass (B) in NM-Rs and mice. Red arrow shows a dip around 0.25% BM. (C-D) Age effect on spleen size in NM-R (C) and in mouse (D): n = 32 NM-Rs (combined ssNM-R and lsNM-R) and n = 40 mice. (E-F) Age effect on liver size in NM-R (E) and in mouse (F): n = 28 NM-Rs (combined ssNM-R and lsNM-R) and n = 40 mice; BM: body mass. Spleen mass and liver mass were expressed as %BM. (G) Sex effect on spleen mass in NM-R and in mouse; NM-Rs: n = 13 males (M) and n = 14 females (F); mice n = 22 males (M) and n = 18 females (F). (H) Influence of the colony on spleen mass: NM-Rs from 3 colonies were used in this part of the study (n = 11 for A, n = 9 for B and n = 7 for C). Linear regression analysis: calculated r square (r^2^) is given for each line. Graphs represent mean ± s.e.m. Unpaired *t* test: p value **<0.01.

**Fig S2:**
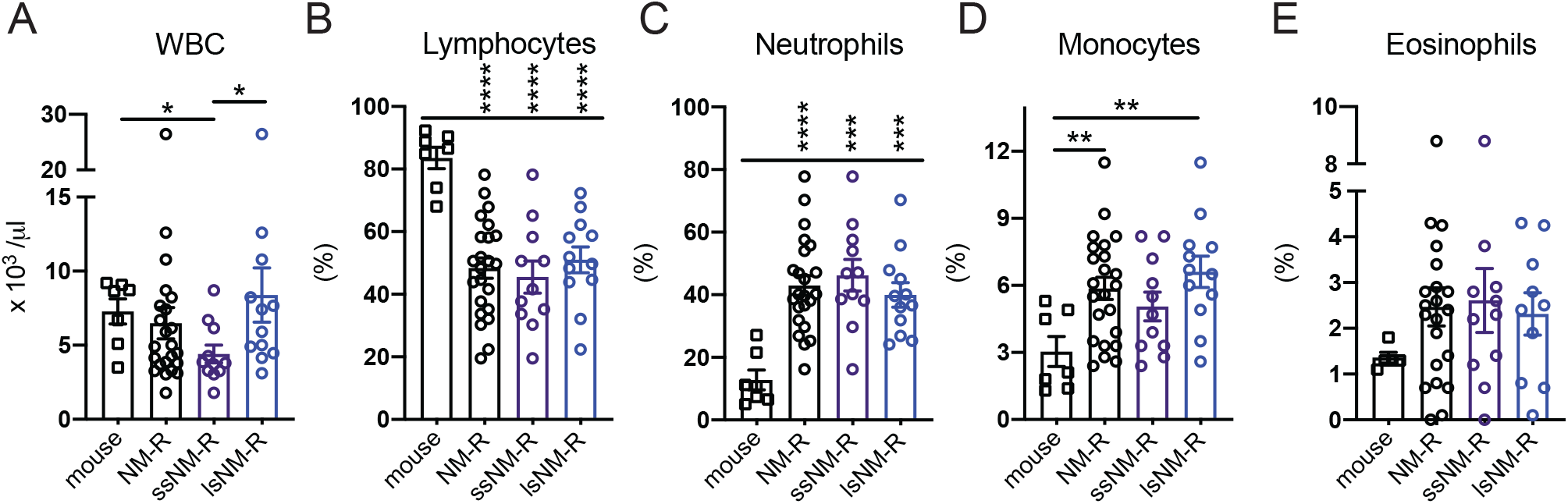
Related to Fig. 2. Peripheral blood count of NM-Rs and mice using a blood cell counter. (A) White blood cell (WBC) count and percentage of (B) lymphocytes, (C) neutrophils, (D) monocytes and (E) eosinophils in the peripheral blood of NM-R and mouse measured using a blood cell counter: n = 11 ssNM-R, n= 12 lsNM-R, n = 23 NM-R (combined ssNM-R and lsNM-R) and n = 7 mice except in (E) n = 11 ssNM-R, n = 10 lsNM-R, n = 21 NM-R (combined ssNM-R and lsNM-R) and n = 5 mice. Graphs represent mean ± s.e.m. Unpaired *t* test: p value *<0.05, **<0.01, ***<0.001 and ****<0.0001.

**Fig S3:**
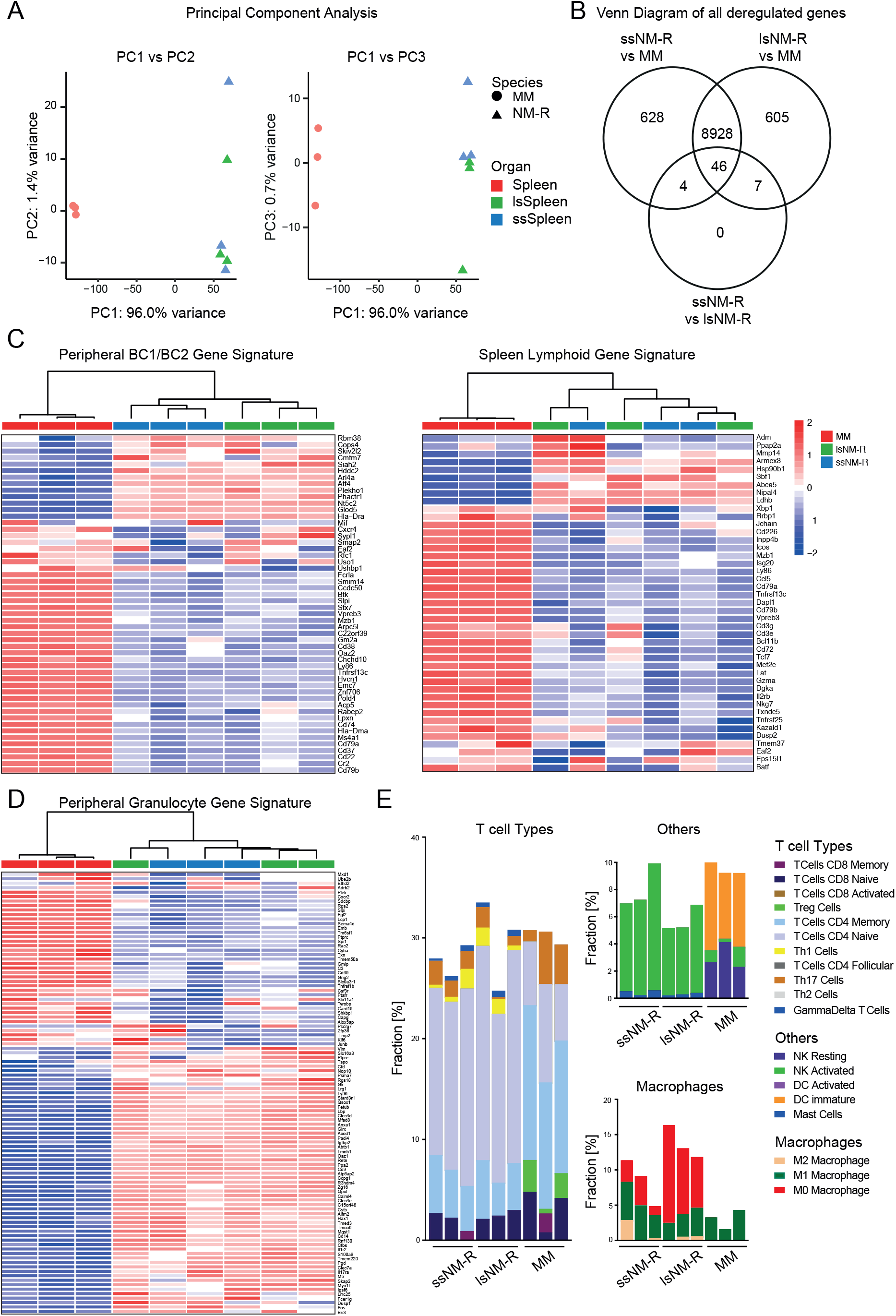
Related to Fig. 3. Transcriptomic analysis of NM-R and mouse spleen. (A) Principal component analysis (PCA) based on the top 3000 most variably expressed genes. Principal component (PC): 1 versus 2 (left panel) and 1 versus 3 (right panel) are displayed. (B) Venn diagram of differentially regulated genes in the spleen of ssNM-R, lsNM-R compared to mouse or to each other. (C) Heatmap of peripheral BC1/BC2 gene sets associated with B cells from [43] (left panel) and of lymphoid gene sets from [16] (right panel) for the spleen of mouse, ssNM-R and lsNM-R. (D) Heatmap of peripheral granulocyte genes from [43] for the spleen of mouse, ssNM-R and lsNM-R. (E) CIBERSORT prediction of fraction of T cell types, macrophage types (M1, M2 and M0) and other cell types (DC: dendritic cells, mast cells and NK: natural killer cells) found in the spleen of ssNM-R, lsNM-R in comparison to mouse. n = 3 per group. Data is based on RNAseq and bars represent fraction in %. MM: mouse (*Mus musculus*).

**Fig S4:**
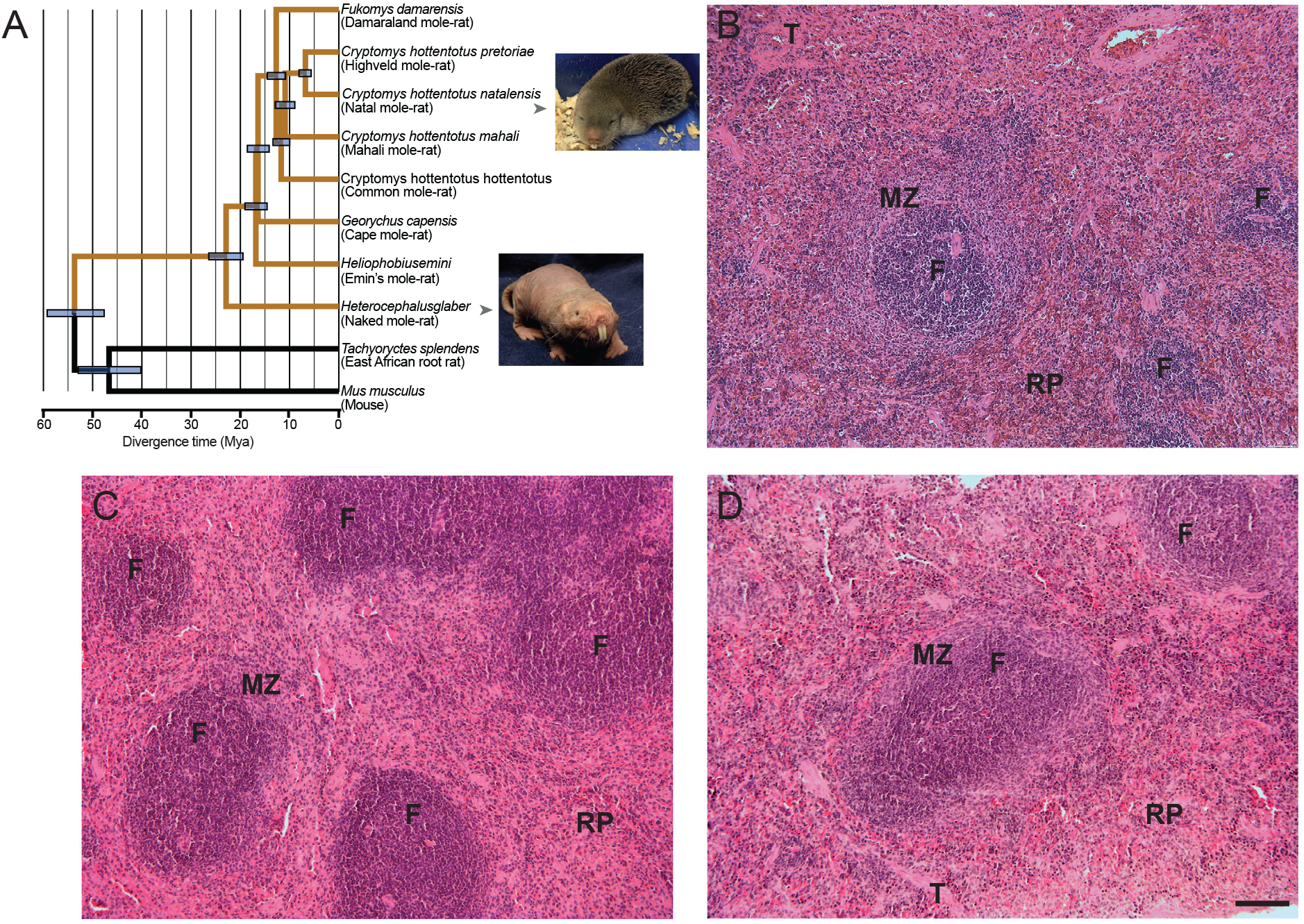
Related to Fig. 4. Spleen morphology of rat, Natal and Highveld mole-rats. (A) Phylogenic tree of known African mole-rats modified from [26]. (B-D) H&E staining of the spleen of rat (B), Natal (C) and Highveld (D) mole-rats showing the presence of follicles (F) and a classical red pulp/white pulp ratio for rodents. MZ: marginal zone, RP: red pulp, T: trabeculae. Scale bar = 100 µm.

**Fig S5:**
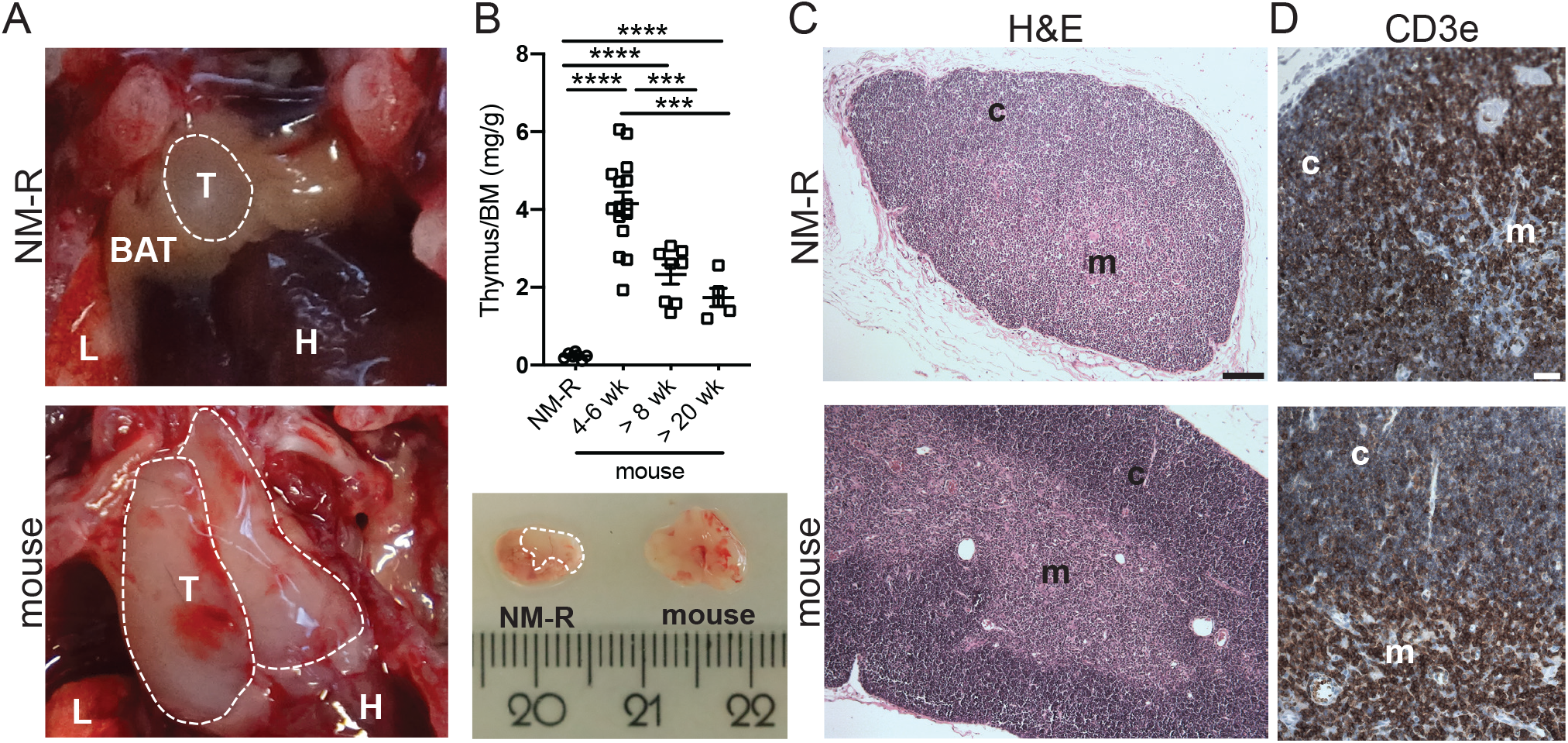
NM-Rs have small thoracic thymus. (A) Representative images of NM-R and mouse thoracic thymus, and (B) their size comparison across mice of different ages and NM-R (top panel): mice n = 15 (4-6 weeks), n = 8 (>8 weeks), n = 4 (>20 weeks) and n = 7 NM-R. Bottom panel: in NM-R the thoracic thymus (white dashed line) is small, and often surrounded by BAT but not in mouse. (C) H&E staining of the thoracic thymus of NM-R and mouse. (D) Immunostaining of the thymic T-cells with CD3e antibody in NM-R and mouse. Data represent mean ± s.e.m. Unpaired *t* test: p value ***<0.001 and ****<0.0001. Scale bars = 100 µm (C), 30 µm (D). T: thoracic thymus, H: heart, BAT: brown adipose tissue, L: lung, c: thymic cortex, m: thymic medulla. BM: body mass in g, thoracic thymus weight in mg.

**Fig S6:**
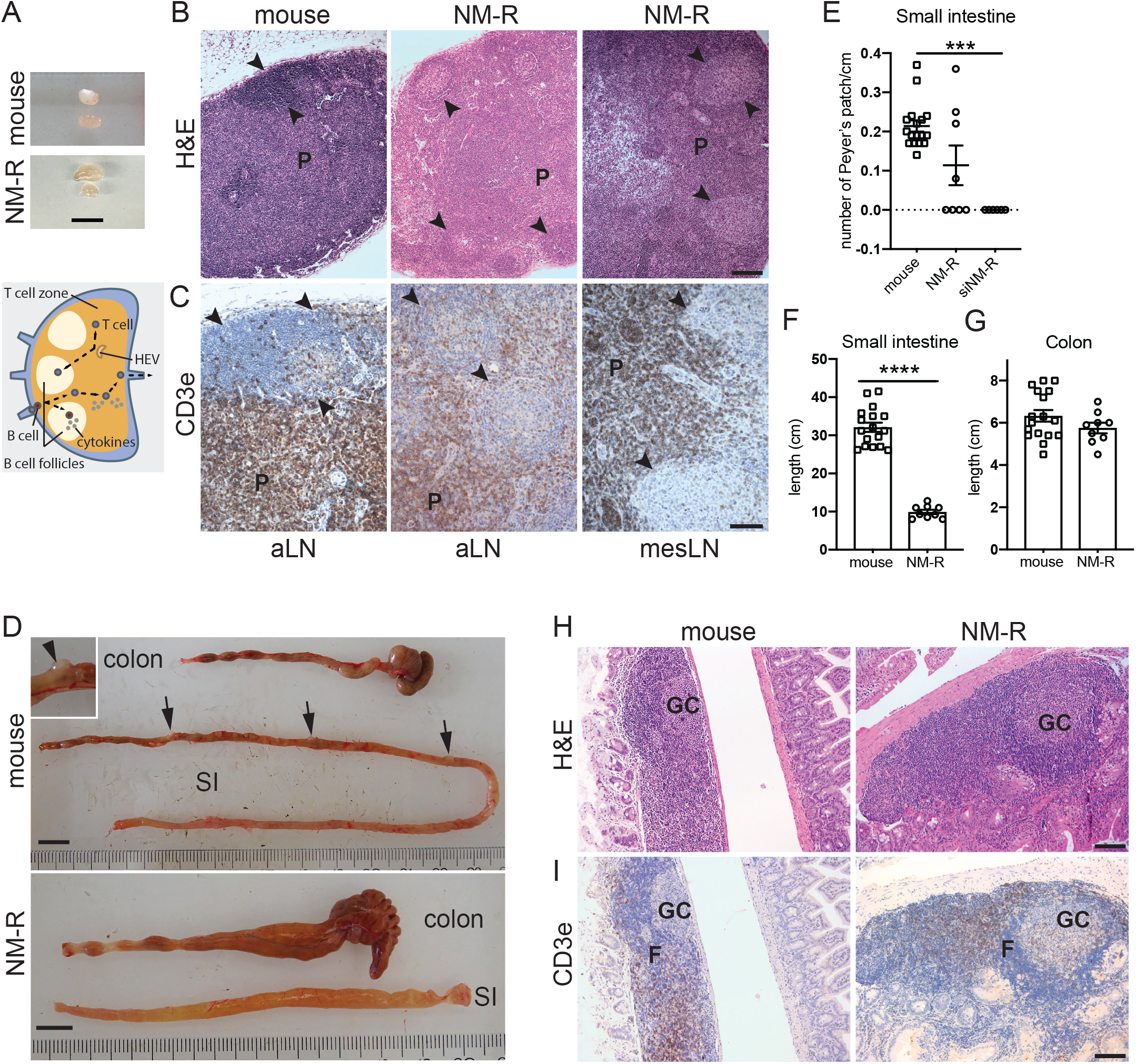
NM-R lymphoid nodules show typical B and T cell areas. (A) Representative image of axillary lymph node (aLN) from NM-R and mouse (top panel) and schematic representation of a lymph node showing the B cells follicles, the T-cell zone, high endothelial venules (HEV) and the migration of lymphocytes (dashed lines) directed by cytokines (bottom panel). (B) H&E staining of aLN and mesenteric lymph node (mesLN) of NM-R and mouse. (C) Immunostaining of T cells in LN with CD3e antibody (brown labelled-cells) in NM-R and mouse. (D) Representative images of the colon and small intestine (SI) of NM-R and mouse showing Peyer’s patches (arrows and inset) in the mouse SI but not in the NM-R SI. (E) Number of Peyer’s patches/cm of small intestine in NM-Rs (n= 8), siNM-Rs (n = 6) and mice (n= 17). (F-G) Length of small intestine (F) and colon (G) in NM-Rs (n = 9) and mice (n = 17). (H) H&E staining of small intestine lymphoid nodule from NM-R and mouse. (I) Immunostaining of T cells with CD3e antibody (brown labelled cells) in the small intestine lymphoid nodule from NM-R and mouse. Note that CD3e+ T cells are found in T cell and B cell zones of aLN, mesLN and small intestine lymphoid nodule from NM-R and mouse (C and I). Follicles: F, germinal center (GC or arrows), P: paracortex. Graphs represent mean ± s.e.m. One-way ANOVA with Tukey’s post-hoc test for multiple comparisons in E; Unpaired *t* test in F and G: p value ***<0.001 and ****<0.0001. Scale bars = 5 mm (A), 50 µm (B, H and I), 30 µm (C) and 1cm (D).

**Fig S7:**
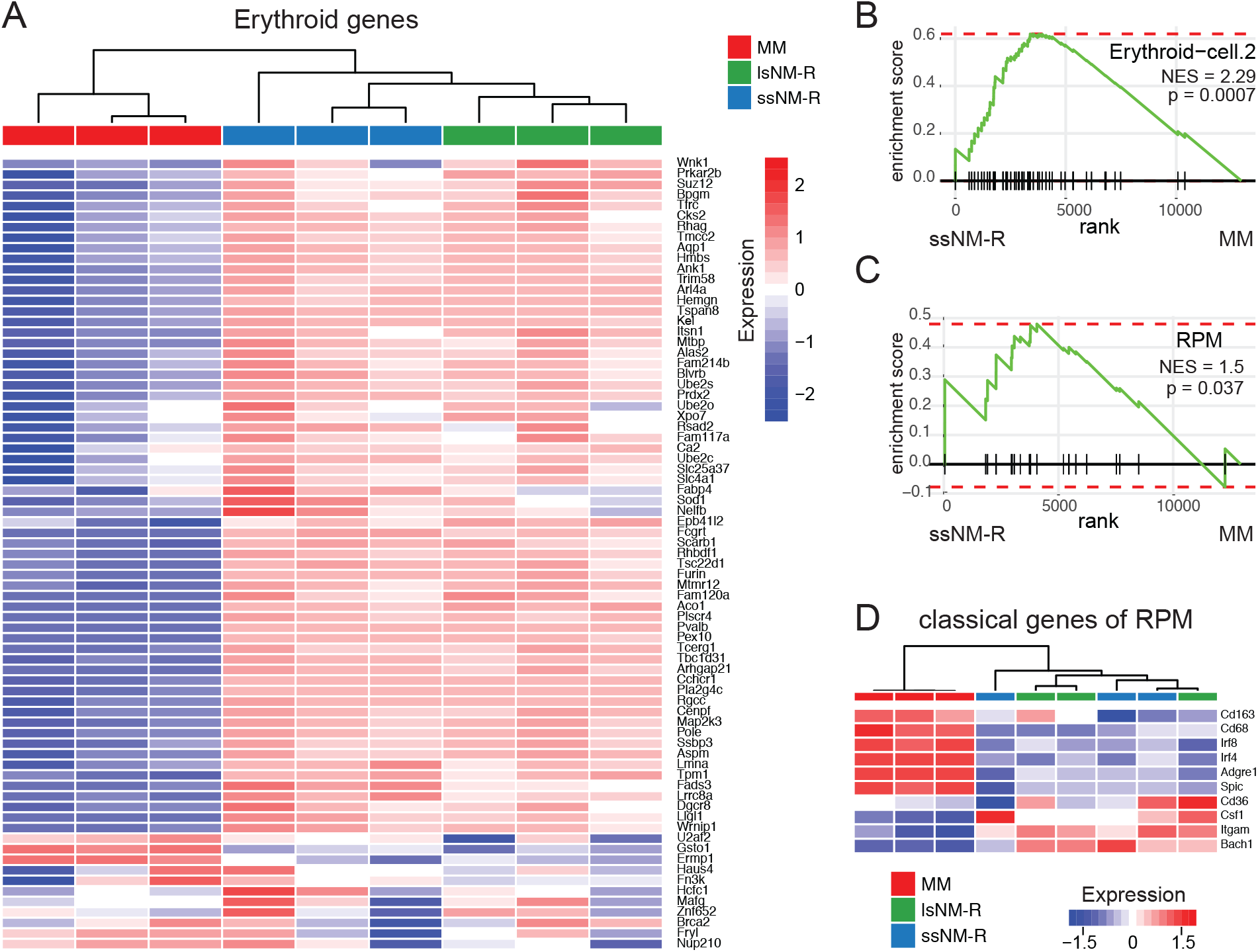
Related to Fig. 6. Extramedullary erythropoiesis occurs in the spleen of ssNM-R and lsNM-Rs. (A) Heatmap of erythroid genes in the spleen of mouse, ssNM-R and lsNM-R. (B) GSEA of erythroid gene sets from [16] and (C) of splenic red pulp macrophage (RPM) gene sets for ssNM-R spleen compared to mouse. (D) Heatmap of classical genes of mouse RPM that are mostly downregulated in NM-R. For all RNA expression, heatmap and GSEA: n =3 mice, n =3 ssNM-Rs and n =3 lsNM-Rs; MM: mouse (Mus musculus), NES: normalized enrichment score; p value is adjusted p value.

**Fig S8:**
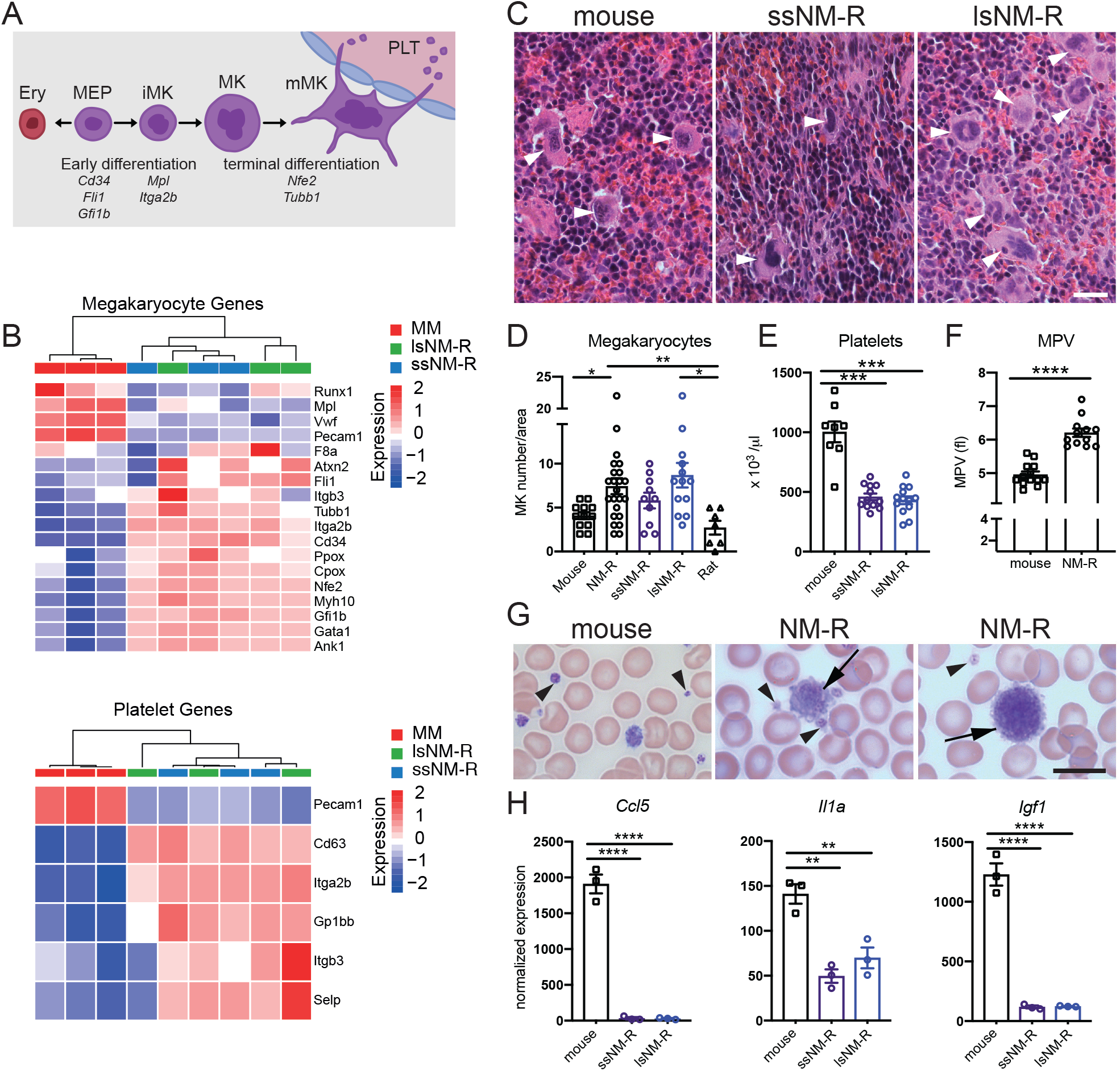
Extramedullary megakaryopoiesis does not contribute to the hyperplasic spleen of lsNM-Rs. (A) Schematic representation of megakaryocytic differentiation. Ery: erythrocyte, MEP: megakaryocyte-erythroid progenitor, iMK: immature MK, mMK: mature MK, PLT: platelets. (B) Heatmap of genes associated with megakaryopoiesis (top) and platelets (bottom) in mouse and NM-R spleen. Samples are hierarchically clustered based on Pearson correlation. (C) H&E staining of megakaryocytes (white arrowheads) in the spleen of mouse, ssNM-R and lsNM-R. (D) Quantification of megakaryocytes (MK) found in mice (n = 12), NM-R (combined ssNM-R and lsNM-R, n = 23), ssNM-R (n = 10), lsNM-R (n = 13) and rat (n = 7). (E) Platelet count in the PB of mice (n = 8), ssNM-R (n = 12) and lsNM-R (n = 13). (F) Mean platelet volume (MPV) in PB of mice (n = 13) and NM-Rs (n = 12). (G) Representative May-Grünwald staining of platelets found in PB smears of mouse and NM-R: mature platelets (arrowhead); immature platelets (arrow). (H) Normalized RNA expression levels of 3 regulatory genes of MK terminal differentiation. Unpaired *t* test in D-F and one-way ANOVA with Tukey’s post-hoc test for multiple comparisons in H: p value *<0.05, **<0.01, ***<0.001 and ****<0.0001. Transcriptomic data is based on RNAseq, MM: mouse (Mus musculus). Data represent mean ± s.e.m, scale bars = 20 μm (C) and 10 μ (G).

**Fig S9:**
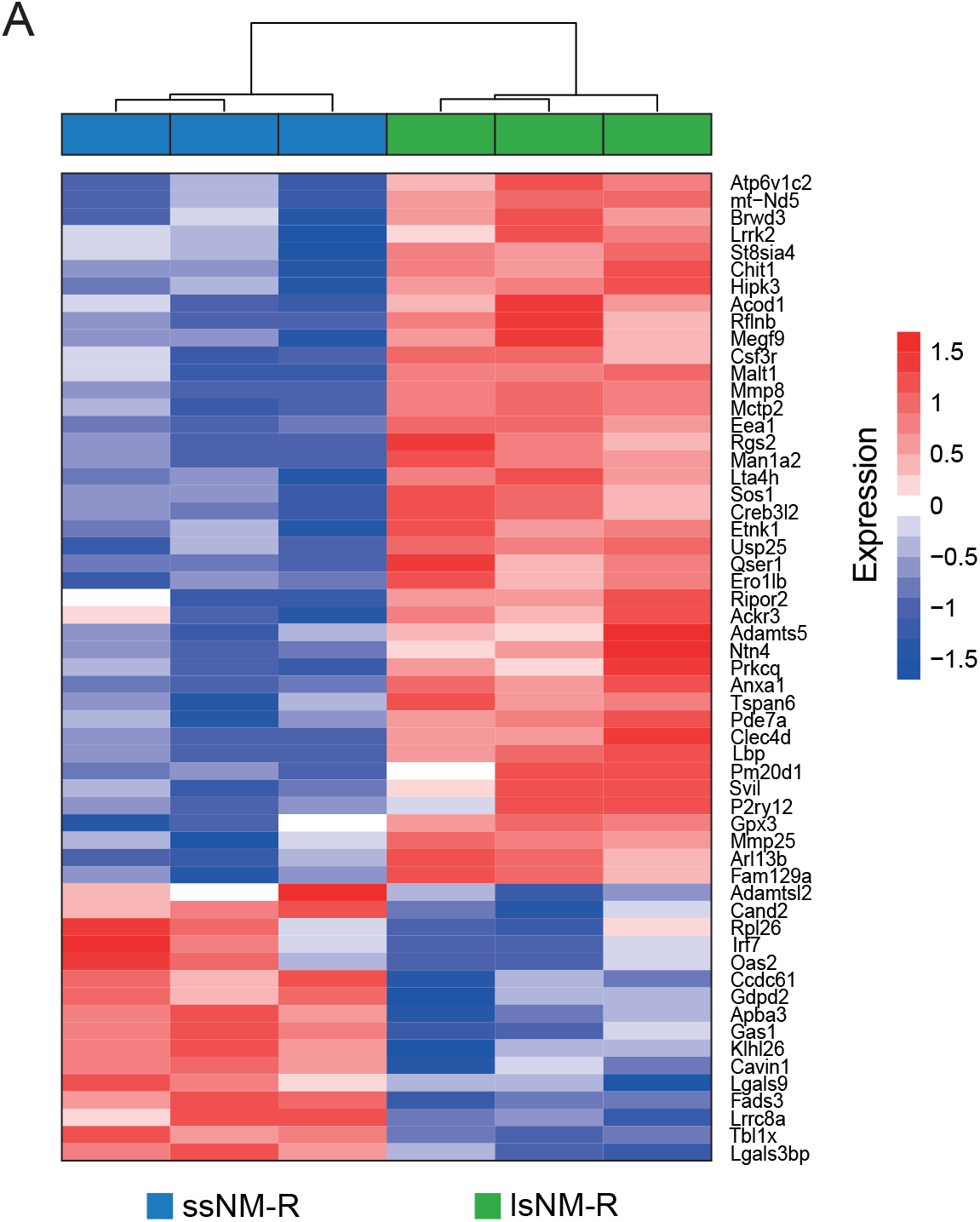
Related to Fig. 7. Comparison of differentially expressed genes between ssNM-R and lsNM-R spleens. Heatmap of all differentially expressed genes in the spleen of lsNM-R in comparison to ssNM-R. Samples are hierarchically clustered based on Pearson correlation. n = 3 per group.

**Table S4:**
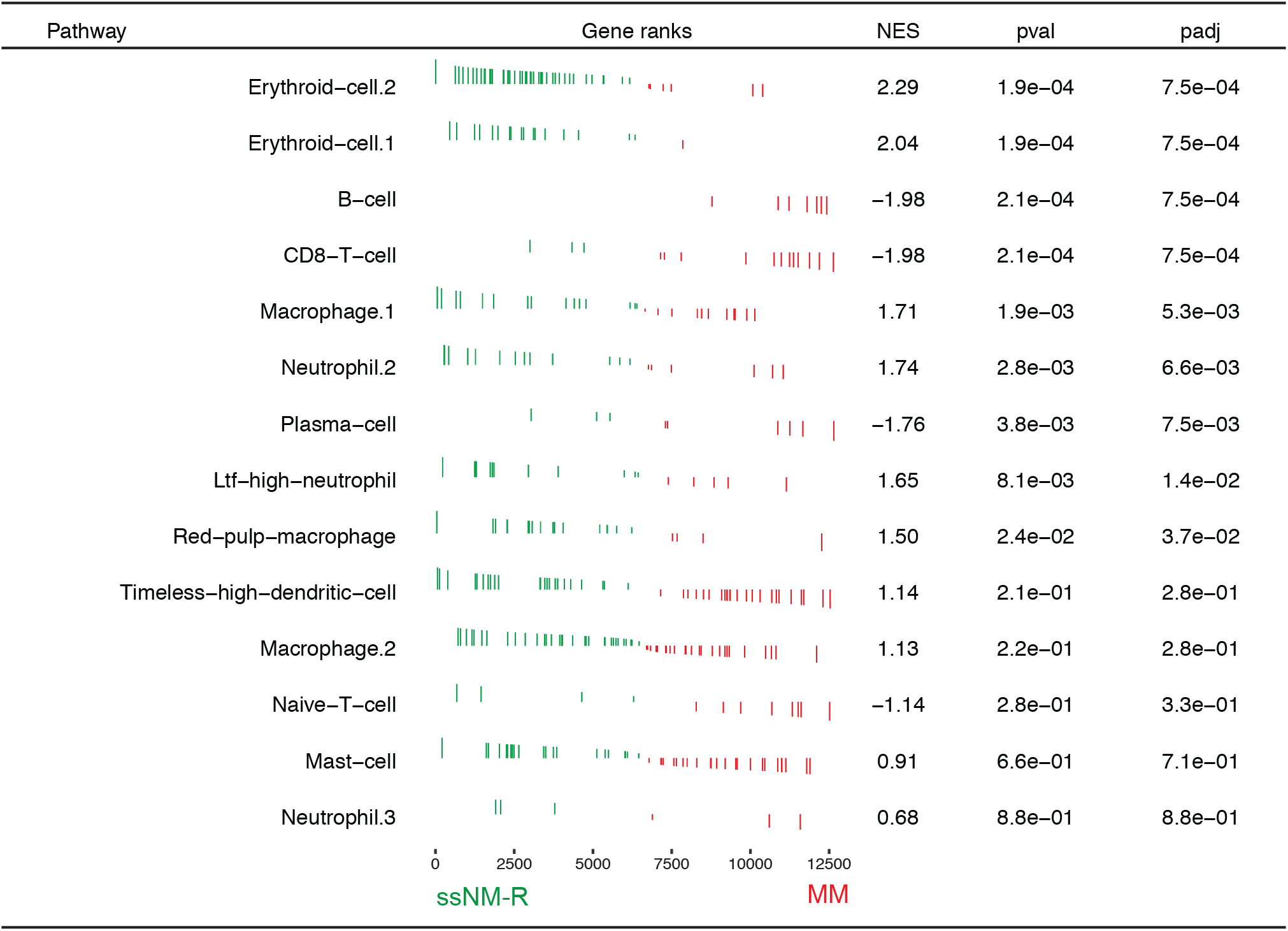
Enrichment analysis of NM-R immune cell gene signatures. GSEA results with custom pathway signature defined by the immune cell signatures from single cell transcriptomic data as described in [16]. Transcriptome data from spleens of ssNM-R versus MM were compared. NES, normalised enrichment score; pval, p value; padj, adjusted p value. For more details, see methods section.

| Pathway | Gene ranks | NES | pval | padj |
| --- | --- | --- | --- | --- |
| Erythroid-cell.2 |  | 2.29 | 1.9e-04 | 7.5e-04 |
| Erythroid-cell.1 |  | 2.04 | 1.9e-04 | 7.5e-04 |
| B-cell |  | -1.98 | 2.1e-04 | 7.5e-04 |
| CD8-T-cell |  | -1.98 | 2.1e-04 | 7.5e-04 |
| Macrophage.1 |  | 1.71 | 1.9e-03 | 5.3e-03 |
| Neutrophil.2 |  | 1.74 | 2.8e-03 | 6.6e-03 |
| Plasma-cell |  | -1.76 | 3.8e-03 | 7.5e-03 |
| Ltf-high-neutrophil |  | 1.65 | 8.1e-03 | 1.4e-02 |
| Red-pulp-macrophage |  | 1.50 | 2.4e-02 | 3.7e-02 |
| Timeless-high-dendritic-cell |  | 1.14 | 2.1e-01 | 2.8e-01 |
| Macrophage.2 |  | 1.13 | 2.2e-01 | 2.8e-01 |
| Naive-T-cell |  | -1.14 | 2.8e-01 | 3.3e-01 |
| Mast-cell |  | 0.91 | 6.6e-01 | 7.1e-01 |
| Neutrophil.3 |  | 0.68 | 8.8e-01 | 8.8e-01 |

